# *Drosophila* insulin-like peptide 1 (DILP1) promotes organismal growth and catabolic energy metabolism during the non-feeding pupal stage

**DOI:** 10.1101/421669

**Authors:** Sifang Liao, Stephanie Post, Philipp Lehmann, Jan A. Veenstra, Marc Tatar, Dick R. Nässel

## Abstract

The insulin/IGF-signaling pathway is central in control of nutrient-dependent growth during development, and in adult physiology and longevity. Eight insulin-like peptides (DILP1-8) have been identified in *Drosophila* and several of these are known to regulate growth, metabolism, reproduction, stress responses and lifespan. However, the functional role of DILP1 is far from understood. Previous work has shown that *dilp1*/DILP1 is transiently expressed mainly during the non-feeding pupal stage and the first days of adult life. Here we show that mutation of *dilp1* diminishes organismal weight during pupal development, whereas overexpression increases it, similar to *dilp6* manipulations. No growth effects of *dilp1* or *dilp6* manipulations were detected during larval development. We next show that *dilp1* and *dilp6* increase metabolic rate in the late pupa and promote lipids as the primary source of catabolic energy. This lipid mobilization in the pupa is not correlated with transcriptional changes of adipokinetic hormone. The effects of *dilp1* manipulations carry over to the adult fly. In newly eclosed flies, survival during starvation is strongly diminished in *dilp1* mutants, but not in *dilp2* and *dilp1*-*dilp2* double mutants, whereas in older flies only double mutants display reduced starvation resistance. In conclusion, *dilp1* and *dilp6* promote growth of adult tissues during the non-feeding pupal stage, likely by utilization of stored lipids. This results in larger newly-eclosed flies with reduced stores of pupal-derived nutrients and diminished starvation tolerance and fecundity.

## Introduction

The Insulin/IGF signaling (IIS) pathway plays a central role in nutrient-dependent growth control during development, as well as in adult physiology and aging [1-5]. More specifically, in mammals insulin, IGFs and relaxins act on different types of receptors to regulate metabolism, growth and reproduction [6-9]. This class of peptide hormones has been well conserved over evolution and therefore the genetically tractable fly *Drosophila* is an attractive model system for investigating IIS mechanisms [4,10,11]. Eight insulin-like peptides (DILP1-8), each encoded on a separate gene, have been identified in *Drosophila* [10,12-14]. The genes encoding these DILPs display differential temporal and tissue-specific expression profiles, suggesting that they have different functions [12,14-17]. Specifically, DILP1, 2, 3 and 5 are mainly expressed in median neurosecretory cells located in the dorsal midline of the brain, designated insulin-producing cells (IPCs) [12,16,18-20]. The IPC derived DILPs can be released into the open circulation from axon terminations in the corpora cardiaca, the anterior aorta and the crop. Genetic ablation of the IPCs reduces growth and alters metabolism, and results in increased resistance to several forms of stress and prolongs lifespan [18,21].

The functions of the individual DILPs produced by the IPCs may vary depending on the stage of the *Drosophila* life cycle. Already the temporal expression patterns hint that DILP1-3 and 5 play different roles during development. Thus, whereas DILP2 and 5 are relatively highly expressed during larval and adult stages, DILP1 and 6 are almost exclusively expressed during pupal stages under normal conditions [15,22].

DILP1 is unique among the IPC-produced peptides since it can be detected primarily during the non-feeding pupal stage and the first few days of adult life when residual larval/pupal fat body is present [15,16]. Furthermore, in female flies kept in adult reproductive diapause, where feeding is strongly reduced, *dilp1*/DILP1 expression is also high [16]. Its temporal expression profile resembles that of DILP6 although this peptide is primarily produced by the fat body, not IPCs [15,22]. Since DILP6 was shown to regulate growth of adult tissues during pupal development [15,22], we asked whether also DILP1 plays a role in growth control. It is known that overexpression of several of the DILPs is sufficient to increase body growth through an increase in cell size and cell number, and especially DILP2 produces a substantial increase in body weight [12,23,24]. In contrast, not all single *dilp* mutants display a decreased body mass. The *dilp1, dilp2* and *dilp6* single mutants display slightly decreased body weight [10,15,22], whereas the *dilp3, dilp4, dilp5* and *dilp7* single mutants display normal body weight [10]. However, a triple mutation of *dilp2, 3*, and *5* causes a drastically reduced body weight, and a *dilp1–4,5* mutation results in even smaller flies [10,25].

There is a distinction between how DILPs act in growth regulation. DILPs other than DILP1 and 6 promote growth primarily during the feeding larval stages when their expression is high [12,23]. This nutrient dependent growth is relatively well understood and is critical for production of the steroid hormone ecdysone and thereby developmental timing and induction of developmental transitions such as larval molts and pupariation [26-30]. The growth during non-feeding stages, which affects imaginal discs and therefore adult tissues, is far less studied. In this study, we investigate the role of *dilp1*/DILP1 in growth regulation in *Drosophila* in comparison to *dilp6*/DILP6. We found that mutation of *dilp1* diminishes body weight and ectopic *dilp1* expression promotes organismal growth during the non-feeding pupal stage, similar to *dilp6*. Determination of metabolic rate and respiratory quotient as well as TAG levels during late pupal development provides evidence that *dilp1* and *dilp6* increase the metabolic rate and that the associated increased metabolic cost is fueled by increased lipid catabolism. We, however, find no evidence for a role of the lipid mobilizing adipokinetic hormone (AKH) [31-33] in the altered lipid catabolism in pupae.

We also investigated the role of *dilp1* mutation and overexpression on early adult physiology. Interestingly, the newly eclosed *dilp1* mutant flies are less resistant to starvation than controls and *dilp2* mutants. Thus, *dilp1* acts differently from other *dilps* for which it has been shown that reduced signaling increases survival during starvation [21]. The decreased starvation resistance in newly hatched flies after *dilp1* overexpression may be a consequence of diminishment of stored nutrients in the pupa during the increased growth of adult tissues, and thus less residual pupal fat body in newborn flies. Also early egg laying and fecundity are affected by *dilp1*.

Taken together, our data suggest that *dilp1*/DILP1 promotes growth of adult tissues during the non-feeding pupal stage, and that this process mainly utilizes stored lipids to fuel the increased metabolic rate. The effect of this increased metabolic rate in the pupa carries over to affect the metabolism in the young adult fly. We suggest that *dilp1*, similar to *dilp6* [15], ensures that if a larva is exposed to poor nutritional conditions it will after pupariation utilize stored nutrients for growth of adult tissues, rather than keeping these stores for the first days of adult life.

## Results

### Mutation of *dilp1* decreases body weight

Growth in *Drosophila* is in part regulated by several of the DILPs through activation of the canonical IIS/TOR (target of rapamycin) pathway [11,12,28]. It was previously reported that decreased *dilp1* activity reduces adult body weight in *Drosophila*, but it was not investigated at what developmental stage this occurred [10,19]. This is relevant to ask since *dilp1* displays a restricted temporal expression during the *Drosophila* life cycle (see Fig 1A). To analyze growth effects of *dilp1* and possible interactions with its tandem-encoded paralog *dilp2*, we employed recently generated *dilp1, dilp2* and double *dilp1-dilp2* null mutants [34]. The efficacy of these mutants was confirmed by qPCR in stage 8-9 pupae and immunolabeling in one-week-old mated female flies (S1 Fig). It can be noted that in *dilp1* mutant pupae the mRNA levels of *dilp2, dilp3* (not shown) and *dilp6* were not altered, but in *dilp6* mutants the *dilp1* level was upregulated (S1A-C Fig). At the protein level DILP2 but not DILP3 immunofluorescence increased in *dilp1* mutants (S1D-G Fig). These findings suggest only minor compensatory changes in other dilps/DILPs in *dilp1* mutants during the pupal stage.

**Fig. 1.**
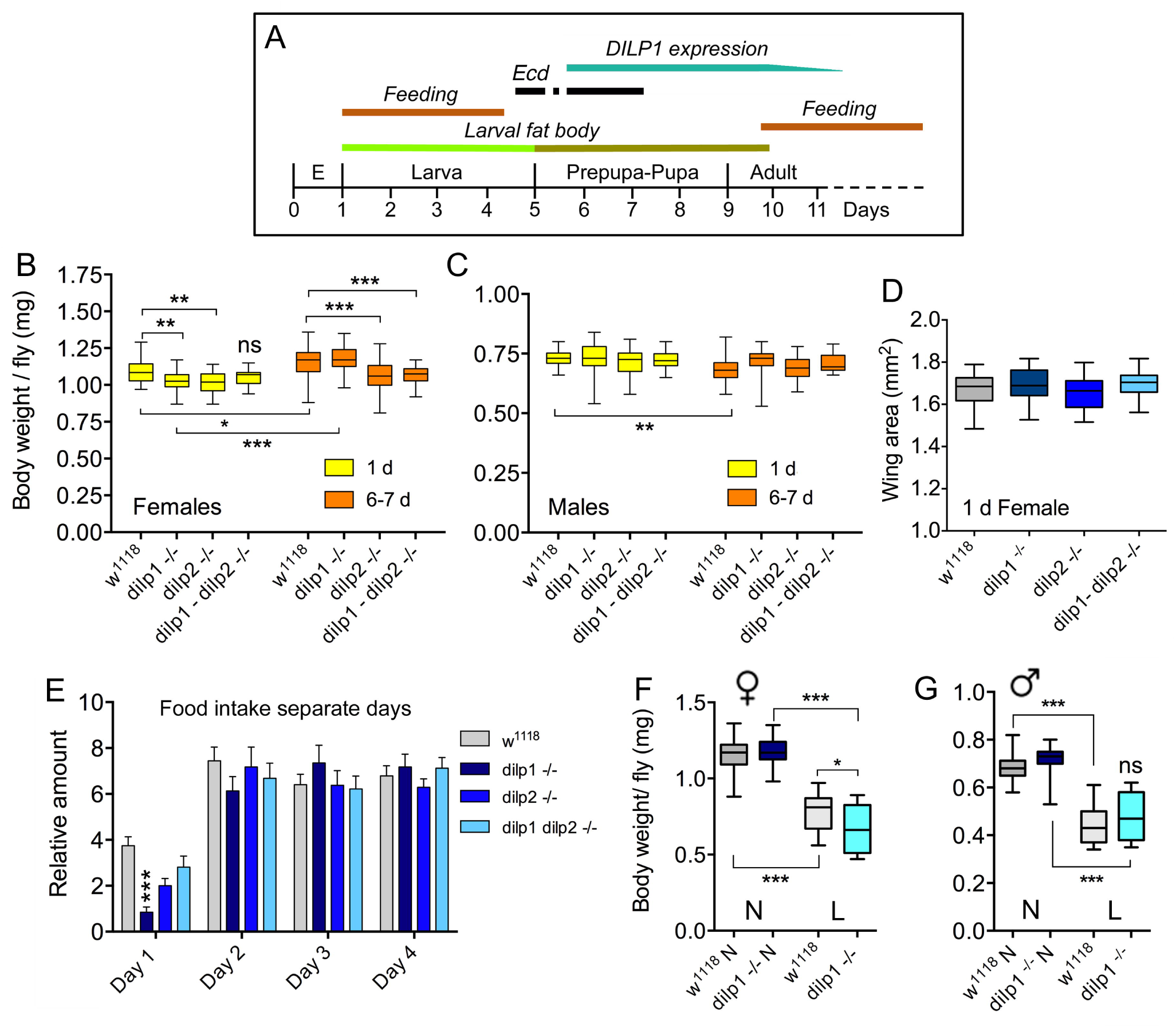
*dilp1* mutant flies display reduced body weight, but are not smaller. **A**. Expression profile of *dilp1*/DILP1 in *Drosophila*. Note that expression of transcript and peptide coincides with the non-feeding pupal stage and the first days of adult life when food intake is reduced (especially day one). It also times with the onset of the second and third ecdysone (*Ecd*) surges in the early pupa (earlier ecdysone peaks are not shown). E, embryo. **B.** Body weight of female flies 1 day and 6-7 days after adult eclosion. *dilp1* mutant flies display reduced body weight when 1 d old, but gain substantially the first week. Also dilp2 mutants weigh less, but do not gain much weight first week. The double mutants are not significantly affected compared to controls at 1 d, but after 6-7 d both *dilp2* and double mutants weigh less that controls and *dilp1* mutants. Data are presented as medians ± range, n = 25–30 flies for each genotype from three independent replicates (*p < 0.05, **p < 0.01, ***p < 0.001, two-way ANOVA followed by Tukey’s test). **C.** In male flies the three mutants display weights similar to controls and controls lose weight the first week. Data are presented as medians ± range, n = 18–30 flies for each genotype from three independent replicates (**p < 0.01, two-way ANOVA followed with Tukey’s test). **D.** Wing area was used as a proxy for organismal growth. The three mutants did not display altered wing size. Data are presented as medians ± range, n = 16–23 flies for each genotype from three independent replicates (One-way ANOVA followed with Tukey’s test). **E**. Food intake was monitored over four days in a CAFE assay. The first day the *dilp1* mutant flies feed less than the other genotypes, whereas during the following days there is no difference between genotypes. Data are presented as means ± S.E.M, n = 20–30 flies for each genotype from three independent replicates (***p < 0.001, two-way ANOVA followed with Tukey’s test). **F** and **G.** Body weight of 7 d old flies that had been exposed to normal diet (N) or low protein diet (L) during late larval stage. The female *dilp1* mutant flies displayed lower body weight than controls after low protein. Data are presented as medians ± range, n = 17-29 flies for each genotype from three replicates (*p < 0.05, ***p < 0.001, one-way ANOVA followed by Tukey’s test).

We monitored the body weight (wet weight) of *dilp1, dilp2* and *dilp1/dilp2* double mutants. First we measured the body weight both in newborn and 6-7 day old adult mated *dilp1* mutant flies. In female flies the newly hatched *dilp1* mutants displayed a decrease in body weight compared to controls (Fig 1B). However, this difference in body weight was no longer detectable in 6-7-day-old mated flies kept under normal feeding conditions; a significant weight increase was observed (Fig 1B). Also *dilp2* mutant female flies have significantly lower body weight than controls one day after emergence, but in contrast to *dilp1* mutants they did not increase the weight over 6-7 days of feeding (Fig 1B). Interestingly the weight of *dilp1/dilp2* double mutants was not significantly affected compared to the single mutants (and control) and no weight increase was seen the first week, except in control flies (Fig 1B). Thus, there was no additive effect of the two mutations. In male flies none of the mutant flies displayed altered body weight (Fig 1C). To determine whether decreased organismal growth was responsible for the lower body weight we measured wing size in the female mutant flies and found no significant difference to controls (Fig 1D). Thus, the decreased weight of the flies does not seem to reflect a significant decrease in organismal size.

We next asked whether the weight gain over the first 6-7 days seen in Fig 1B was caused by increased feeding. Using a capillary feeding (CAFE) assay over four days, we found that during the first day of assay the *dilp1* mutant flies actually fed less than the other mutants and control flies (Fig 1E). The subsequent days food intake was not significantly different between the genotypes. Thus, the food intake profile does not explain the weight gain over the 6-7 days (Fig 1E); possibly the female *dilp1-/-* flies excrete less waste or spend less energy. It was shown earlier that 1 week old *dilp1* mutant flies display a two-fold increased expression of *dilp6* transcript [16], that might compensate for the loss of *dilp1*. However, in the midpupal stage there is no significant upregulation of *dilp6* in *dilp1* mutants (S1C Fig).

In a study of *dilp6* it was shown that if third instar larvae (after reaching critical size) were put on a low protein diet, they emerged as smaller adults and that this was accentuated in *dilp6* mutants [15]. This suggests that *dilp6* is important for assuring growth of adult tissues under low protein conditions. We, thus, performed a similar experiment with *dilp1* mutant larvae kept on normal food or low protein diet. Flies emerging from larvae on restricted protein indeed displayed significantly lower body weight and female *dilp1* mutants weighed less than controls under protein starvation (Fig 1F). In male flies this latter effect was not seen in the mutants (Fig 1G).

We then asked whether mutation of both *dilp1* and *dilp6* would result in a further decrease of body weight and generated a recombinant *dilp1-dilp6* mutant. Using qPCR we found that these flies displayed virtually no detectable *dilp1* and *dilp6* RNA (S2A Fig.). The weights of *dilp1/dilp6* mutants were significantly reduced compared to controls (S2B Fig.). However, their weights were not diminished more that those of the single *dilp1* and *dilp6* mutants, indicating that there was no additive effect of loss of both *dilps*.

### Overexpression of *dilp1* promotes growth during the non-feeding pupal stage

Having shown effects of the *dilp1* null mutation on adult flyweight we next explored the outcome of over-expressing *dilp1*, either in IPCs, or more broadly. For this we generated several UAS-*dilp1* lines [see [34]]. These UAS-*dilp1* lines were verified by DILP1 immunolabeling after expression with several Gal4 drivers (S3A-D Fig) and by qPCR in stage 8-9 pupae (S4A-F Fig). Overexpression of *dilp1* in fat body (*ppl*-Gal4 and *to*-Gal4) and IPCs (*dilp2*-Gal4) results in a drastic upregulation of *dilp1* RNA (S4A, D Fig), but has no effect on *dilp2* and *dilp6* expression (S4B, C, E, F Fig), except a minor decrease in *dilp2* for *ppl*-Gal4 (S4B Fig). At the protein level *dilp1* overexpression resulted in minor changes in DILP2, 3 and 5 immunolevels in IPCs of one week old adult female flies (S5A-E Fig). One line, UAS-*dilp1* (III), was selected for subsequent experiments since it generated the strongest DILP1 immunolabeling.

First, we used a *dilp2*-Gal4 driver to express *dilp1* in the IPCs and detected a significant increase in body weight of female flies (Fig 2A). We then expressed *dilp1* in the fat body, the insect functional analog of the liver and white adipocytes in mammals [35-37]. The fat body displays nutrient sensing capacity, and is an important tissue for regulation of growth and metabolism in *Drosophila* [15,37-41]. It is also the tissue where DILP6 is produced and released [15,38]. To investigate the effect of ectopic *dilp1* expression in the fat body, we used the fat body-specific *pumpless (ppl)* and *takeout (to)* Gal4 drivers. The efficiency of the drivers was confirmed by DILP1 immunostaining of larval fat body of *ppl>dilp1* and *to>dilp1* flies, but not in the control flies (S3D Fig). In *ppl>dilp1* flies we also found DILP1 labeling in the nephrocytes (not shown), which are highly endocytotic cells located close to the heart [42]. The immunoreactive DILP1 is likely to have accumulated from the circulation after release from the fat body since the *ppl*-Gal4 is not expressed in the nephrocytes.

**Fig. 2.**
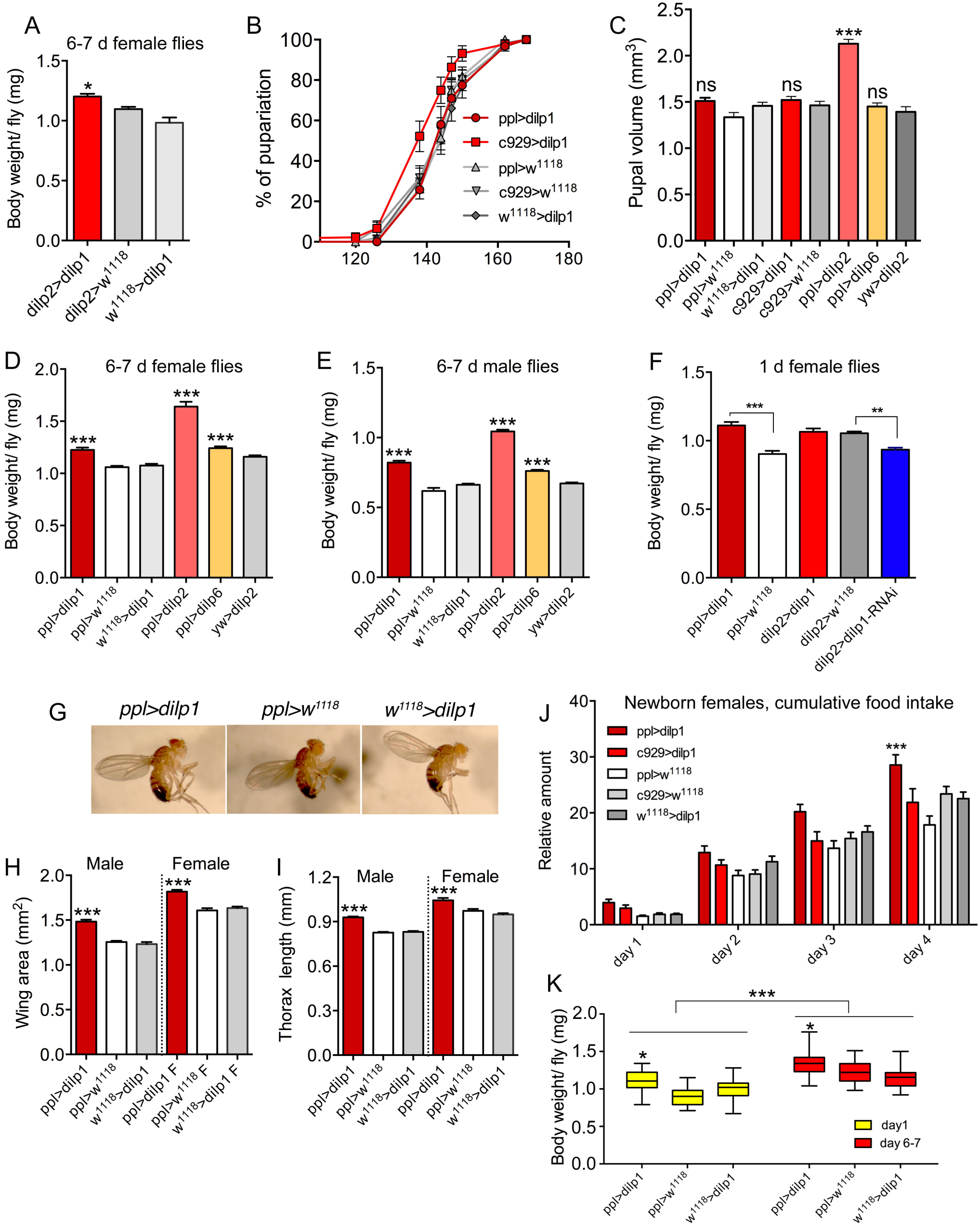
Overexpression of *dilp1* affects growth during pupal stage. **A.** Expression of *dilp1* in insulin-producing cells (IPCs) with *dilp2*-Gal4 driver increases body weight of 6-7 d adult flies. Data are presented as means ± S.E.M, n = 14–23 flies for each genotype from three independent replicates (*p < 0.05, one-way ANOVA followed by Tukey’s test). **B.** Overexpression of *dilp1* in fat body (*ppl*-Gal4) or neuroendocrine cells (*c929*-Gal4) does not affect time to pupariation (larval development). Data are presented as means ± S.E.M, n = 138-147 flies for each genotype from three independent replicates (*p < 0.05, as assessed by Log-rank (Mantel-Cox) test). **C.** Overexpression of *dilp1* using *ppl*-Gal4 or *c929*-Gal4 does not affect pupal volume (proxy for larval growth). Also *dilp6* overexpression has no effect, whereas *dilp2* expression triggers a significant increase in pupal volume. Data are presented as means ± S.E.M, n = 15–32 flies for each genotype from three independent replicates. (***p < 0.001, one-way ANOVA followed with Tukey’s test). **D** and **E**. Overexpression of *dilp1, dilp2* and *dilp6* in fat body all lead to adult flies (one week old) with increased body weight both in males and females. Data are presented as means ± S.E.M, n = 24-30 flies for each genotype from three independent replicates. Except for ppl>dilp2, 13 flies were used (*p < 0.05, one-way ANOVA followed with Tukey’s test). **F.** Also one-day-old female flies weigh more than controls after *ppl>dilp1*, but not after *dilp2>dilp1*. Knockdown of *dilp1* by *dilp2*>*dilp1*-RNAi lead to decreased body weight. Data are presented as means ± S.E.M, n = 20-27 flies for each genotype from three independent replicates (**p < 0.01, ***p < 0.001, unpaired Students’ t-test). **G.** Images of flies overexpressing *dilp1* in the fat body and controls. **H** and **I**. Overexpression of *dilp1* in fat body results in flies with increased wing area (H), and length of thorax (I) as proxies for organismal growth. Data are presented as means ± S.E.M, (***p < 0.001, one-way ANOVA followed with Tukey’s test); in H n= 17-24 flies and in I n = 9–17 flies from three independent replicates. **J.** Food intake (CAFE assay) is increased over four days (cumulative data shown) in flies overexpressing *dilp1* in fat body, but not in neuroendocrine cells (c929 Gal4). Data are presented as means ± S.E.M, n = 15–30 flies for each genotype from three independent replicates (*p < 0.05, two-way ANOVA followed with Tukey’s test). **K.** Body weight of 6-7 d female flies is increased for all genotypes compared to 1 d flies. The ppl>dilp1 flies weigh more than controls at both time points. Data are presented as medians ± range, n = 23–27 flies for each genotype from three independent replicates (*p < 0.05, ***p < 0.001, two-way ANOVA followed with Tukey’s test).

Before monitoring the effect of *dilp1* overexpression in the fat body on adult body weight and organismal size, we wanted to determine whether *dilp1* has an effect on larval development. We therefore measured the time to pupariation and size of pupae to determine whether *dilp1* overexpression affected timing of larval development and growth during this stage. Using the *ppl*-Gal4 driver we did not observe any effect on the time from egg to pupa compared to controls (Fig 2B). Pupal volume, as a measurement of larval growth, was not altered by *ppl*-Gal4>*dilp1* (Fig. 2C). As expected [15,38], over-expression of *dilp6* also had no effect on pupal size (Fig 2C). However, as shown earlier for ubiquitously expressed *dilp2* [23], *dilp2* expression in the fat body generated a strong increase in pupal volume, suggesting growth during the larval feeding stage (Fig 2C). Driving *dilp1* with the *c929* Gal4 line, that directs expression to several hundred *dimm*-expressing peptidergic neurons including IPCs [43], we did not observe any effect on time to pupariation or pupal volume (Fig 2B, C). Taken together our data suggest the ectopic *dilp1* does not affect larval growth or developmental time.

Next, we determined the body weight of mated 6-7 d old flies. Body weight increased significantly in *ppl>dilp1* flies compared to the controls both in female (Fig 2D) and male flies (Fig 2E). Here we additionally noted increased weight for *ppl>dilp2* and *ppl>dilp6* flies. We also monitored the weight of one day old flies and found that *ppl>dilp1*, but not *dilp2>dilp1* flies displayed increased weight (Fig 2F). However, *dilp2*>*dilp1*-RNAi induced a decrease in body weight (Fig 2F). Moreover, organismal size, estimated by wing size (Fig 2G, H) and thorax length (Fig 2G, I), increased after ectopic expression of *dilp1* in the fat body. Since we see no effect of *dilp1* expression on developmental time or pupal volume, but register increased body weight and size of adults, we propose that *dilp1*, like *dilp6*, promotes growth of adult tissues during the pupal stage.

It was suggested that *dilp6* promotes growth of adult tissues during pupal development by utilizing nutrients stored in the larval fat body, which is carried into the pupa [15]. This may be the case also for *dilp1*, and if so, newly hatched *dilp1* overexpressing flies should have less energy stores in the form of residual larval fat body. To test this we monitored feeding in recently hatched *dilp1* mutant flies and controls. Indeed, flies overexpressing *dilp1* displayed increased food ingestion over the first four days after adult emergence compared to controls (Fig 2J). Next we compared the weights of one day old and 6-7 day old flies after dilp1 overexpression with *ppl*-Gal4 and found that at both ages the female *ppl>dilp1* flies weighed more than controls and that the older flies were heavier than the younger ones (Fig 2K). In male flies *ppl>dilp1* also increased the body weight, but there was a loss of weight for all genotypes over the first 6-7 days of adult life (S6A Fig). As a comparison *dilp2>dilp1* had only minor effects on body weight of female flies, only in 6-7 d old flies there was an increase (S6B Fig), whereas in males a significant increase was noted at both ages for dilp2>dilp1, and a loss of weight over the next six days for all genotypes (S6C Fig).

Using the *to*-Gal4 fat body driver to express *dilp1* we also noted an increase in weight of recently emerged female and male flies (S6D, E Fig), but no change in body size except a minor increase in thorax length in females (S6F, G Fig). The female *to*>*dilp1* flies increased further in weight the first 6-7 days of adult life, but not later (S6D Fig), whereas the males did not (S6E Fig). Furthermore, with the *to*-Gal4 driver there was no increase in pupal volume, supporting that *dilp1* does not affect larval growth (S6H Fig).

Ectopic expression of *dilp1* in neuroendocrine cells by means of the *c929*-Gal4 increased adult body weight (S7A Fig), but had no effect on wing size in males and females or food intake in young flies (S7B, C Fig), suggesting that *dilp1* expression (and/or systemic release) was not strong enough to yield major effects. Also *dilp2*>*dilp1* flies were tested in food intake and no effect was seen (S7C Fig).

### Overexpression of *dilp1* increases the size of the adult brain and neuroendocrine cells

It was previously shown that signaling through the *Drosophila* insulin receptor (dInR) can lead to an enlargement of cell bodies of neuroendocrine cells in a cell autonomous manner, and that *dilp6* in glial cells is a candidate ligand to mediate this dInR dependent growth [44,45]. Since *dilp1* has a temporal expression profile similar to *dilp6*, and promotes growth of adult tissues in the pupal stage, we asked whether *dilp1* also affects size of neuroendocrine cells that differentiate in the pupa. Thus, we overexpressed *dilp1* with the broad neuroendocrine cell driver *c929*-Gal4 [43,46], and monitored the cell body size of several groups of neuroendocrine cells in the adult CNS with specific peptide antisera. We found that the cell body size of IPCs increased in adult *c929>dilp1* flies, as shown by anti-DILP2 staining (S8A1-3 Fig, Table 1). Furthermore, the cell bodies of the adult-specific pigment-dispersion factor (PDF) expressing clock neurons (l-LN_v_s), as shown here by anti-PDF staining, were enlarged in *c929>dilp1* flies compared to the controls (S8B1-3 Fig, Table 1). Next, we monitored the cell-body size of leucokinin (LK) producing neurons in the abdominal ganglia (ABLKs), and found that the adult-specific anterior, but not the larval-derived posterior ABLKs, displayed increased size in *c929>dilp1* flies (S8C1-3 Fig, Table 1).

**Table 1.** Overexpression of *dilp1* in neurosecretory cells and fat body affects size of brain and certain neurons

| Genotype | Neuron/tissue<sup>1</sup> | Size change | Significance | Figure |
| --- | --- | --- | --- | --- |
| *c929>dilp1* | IPC | + | p<0.001 | S Fig. 7A |
| *c929>dilp1* | I-LNv | + | p<0.01 | S Fig. 7B |
| *c929>dilp1* | ABLK<sub>a</sub> | + | p<0.05 | S Fig. 7C |
| *c929>dilp1* | ABLK<sub>p</sub> | nc | ns | S Fig. 7C |
| *c929>dilp1* | Whole brain | + | p<0.001 | S Fig. 7D |
| *c929>dilp1* | ABLK larva | nc | ns | S Fig. 8E, F |
| *c929>dilp1* | CNS larva | nc | ns | S Fig. 8G |
| *dilp2>dilp1* | IPC | nc | ns | S Fig. 7E |
| *ppl>dilp1* | I-LNv | + | p<0.01 | S Fig. 7G |
| *ppl>dilp1* | Whole brain | + | p<0.05 | S Fig. 7H |
| *pdf>dilp1* | I-LNv | nc | ns | S Fig. 8A, B |
| *pdf>dilp1* | Whole brain | nc | ns | S Fig. 8C |
| *repo>dilp1* | I-LNv | nc | ns | S Fig. 9A, B |

Notes
+, increased size
nc, no change
ns, not significant
<sup>1</sup> Neurons or tissues monitored for size

**Table 2.**
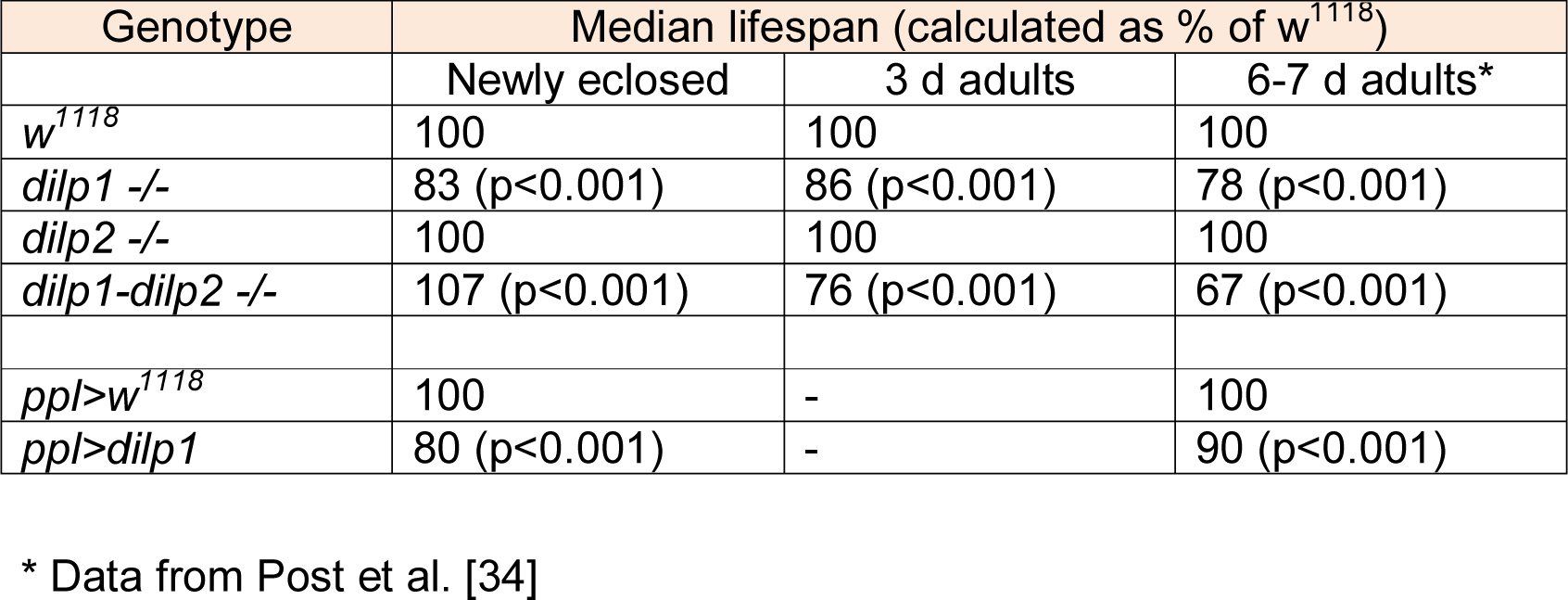
Median lifespans of female flies exposed to starvation.

However, the observed increase in cell body size appears to be partly due to a broader growth of the adult fly tissues, since we found that also the size of the brain increased in *c929>dilp1* flies (S8D Fig, Table 1). The c929-Gal4 is expressed in IPCs and several other groups of peptidergic neurosecretory cells [43,46], which could be the source of systemic release of ectopic DILP1 that affects brain and cell growth. To support that systemic DILP1 is required to promote this growth we employed the *ppl*-Gal4 to drive *dilp1* in the fat body and found an increase in the size of the PDF expressing clock neurons (S8F1-3 Fig, Table 1) and the brain (S8G Fig, Table 1). In contrast, we found that expressing *dilp1* in interneurons, such as PDF-expressing clock neurons does not induce growth of brain neurons (S9A, B Fig, Table 1) or size of the brain (S9C Fig, Table 1), but affected the intensity of PDF immunolabeling (S9D Fig). Thus, paracrine release of DILP1 in the brain does not seem to affect growth of neurons. Interestingly, we found that in third instar larvae, the cell body size of ABLK neurons or the size of the CNS were not different in *c929>dilp1* larvae compared to controls (S9E-G Fig, Table 1), further supporting that *dilp1* overexpression has no effect on cell growth during the larval stage. Finally, since overexpression of *dilp6* in glial cells by *Repo*-Gal4 promotes increase in size of neuronal cell bodies [45], we tested overexpression of *dilp1* in these cells, but found no significant effect on the cell-body size of PDF neurons (S10A, B Fig, Table 1). This again indicates that to affect cell/tissue growth DILP1 must act systemically rather than in a paracrine fashion.

### Metabolic rate and respiratory quotient in pupae of different genotypes

To investigate the role of *dilp1* in utilization of nutrients during pupal development we determined metabolic rate (MR) and respiratory quotient (RQ) in pupae of different genotypes. First we characterized the metabolic trajectory in control pupae (*w*^1118^) by measuring cumulative MR daily throughout pupal development (Fig 3A). These data show the exponential MR curve typical for developing insects, including *D. melanogaster* [47]. To minimize handling stress, we chose to investigate only the end of pupal development in more detail and measured MR and RQ in 4-day-old pupae (that is the cumulative MR between hours 96 and 120 after pupation). For this experiment we used only *ppl*-Ga4 overexpression animals, since the mutant animals displayed high mortality in the respirometry setup used here. As can be seen in Fig 3B and 3C the *ppl>dilp1* and *ppl>dilp6* differed significantly from the controls. The MR was higher and RQ lower in the overexpression flies than in the control flies. RQ values, around 0.6 in both overexpression lines, suggest pure lipid metabolism [48], and lipids are known to be a major or sole fuel during metamorphosis of insects [49,50]. Our findings strongly suggest that *dilp1* (and *dilp6*) affects metabolism in the pupa, maybe to ensure that enough fuel is allocated for growth of adult tissues.

**Fig. 3.**
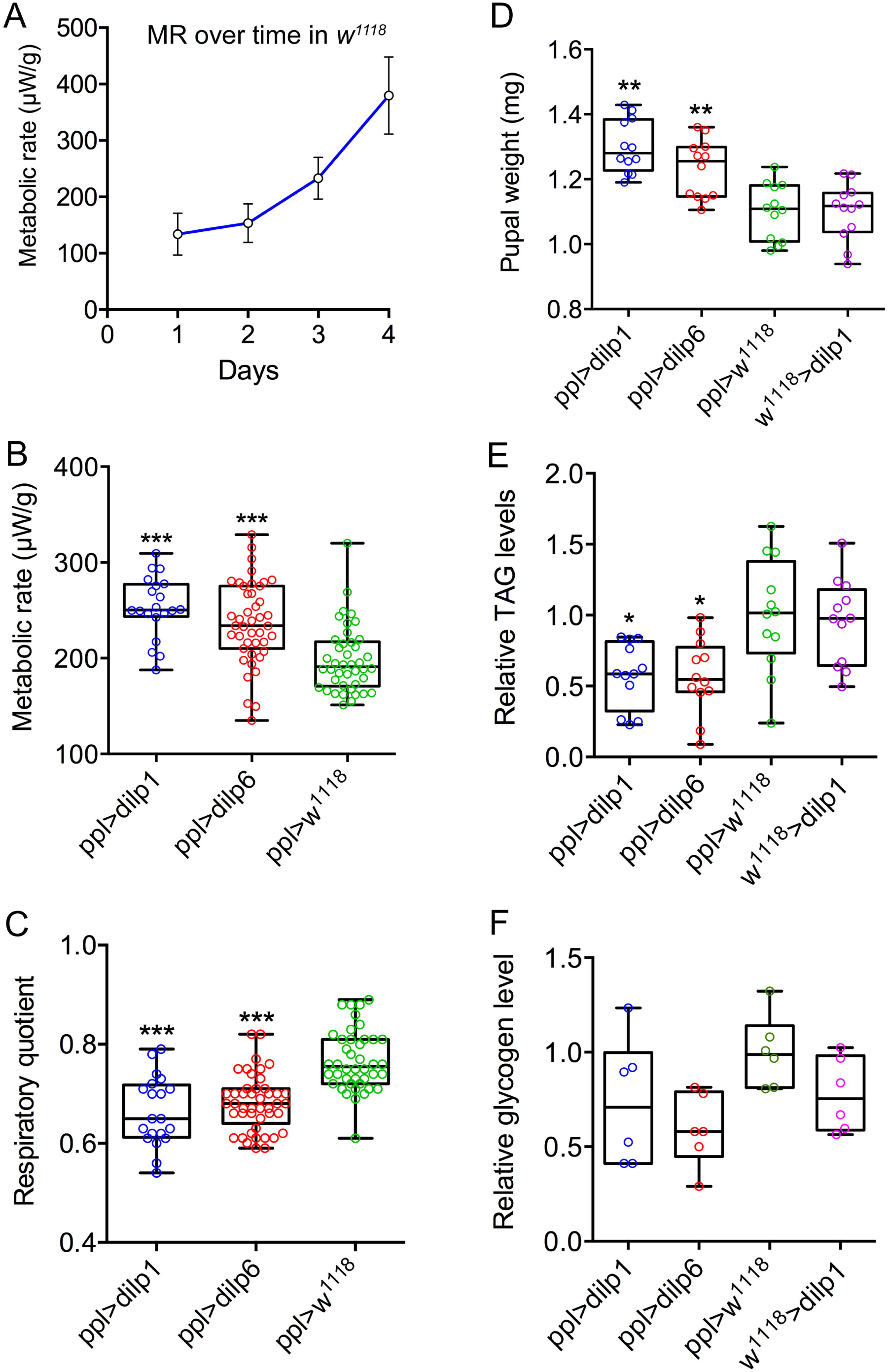
Metabolic rate trajectories and respiratory quotients (RQ) during pupal development respond to *dilp1* and *dilp6* overexpression in the fat body. **A.** Metabolic rate in w^1118^ flies increased exponentially as a function of time. For the ensuing overexpression analysis we studied the period 96-120 hours after pupation. Data are presented as means ± S.E.M, n = 10–15 flies from three independent replicates. **B.** Metabolic rate was significantly elevated during this period in *dilp1* and *dilp6* overexpression flies (*ppl-Gal4*) when compared to *w*^*1118*^ flies. Data are presented as means ± S.E.M, n = 10–15 flies for each genotype from three independent replicates (***p < 0.001, compared to *w*^*1118*^ flies, as assessed by two-way ANOVA followed with Tukey’s test). Data are from both males and females as no difference was found in the ANOVA for sex. **C.** RQ, reflecting catabolic energy substrate, was significantly lower in the overexpression flies when compared to the control flies and indicates a shift from mixed fuel catabolism (RQ = 0.7-0.8) to predominantly lipid catabolism (RQ < 0.7). Data are presented as means ± S.E.M, n = 10–15 flies for each genotype from three independent replicates (***p < 0.001, compared to *w*^*1118*^ flies, as assessed by one-way ANOVA followed with Tukey’s test). Data are from both males and females as no difference was found in the ANOVA for sex. **D.** Four day old pupae (mixed male and female) were weighed (wet weight) before extraction and TAG determination. Overexpression of *dilp1* and *dilp6* both resulted in increased pupal weight. **E**. Levels of TAG were measured in the pupae used for weighing in D. Overexpression of each *dilp* resulted in decreased TAG levels. **F.** Glycogen levels in 4 d old pupae (no significant changes). In **D-E** 12 replicates per genotype with 4 pupae in each replicate (each data point represents 4 pupae), in F 6 replicates per genotype with 4 pupae in each replicate (*p < 0.05, **p < 0.01, one-way ANOVA followed by Tukey’s test).

### TAG, carbohydrates and AKH signaling in pupae of different genotypes

To determine whether it indeed are lipids that fuel growth of adult tissues in 4 day old pupae we determined TAG levels after over expression of *dilp1* and *dilp6* in fat body (*ppl*-Gal4). Pupae of both genotypes displayed increased weight (Fig. 3D) and also significantly reduced TAG levels (Fig. 3E), compared to controls of the same age. The decreased glycogen levels in pupae after ectopic expression of *dilp1* and *dilp6* were not significant (Fig. 3F) and glucose levels were not significantly changed (S11 Fig).

Since AKH is known to mobilize lipids in insects, including *Drosophila* [31-33,51], we determined levels of *akh* transcript in pupae with *dilp1* and *dilp6* overexpression (using *ppl*-Gal4) and in *dilp* mutants, at two different time points (2 and 4 d old pupae). There was no significant alteration in *Akh* transcript after *dilp*1 or *dilp6* overexpression; the only phenotype was a slight upregulation in *dilp1* mutants in 4 d pupae (S11A-D Fig). Next we analyzed levels of transcript of brummer (*bmm*), a lipase known to promote TAG mobilization [52], in pupae of the same stages and found no significant change in expression for any genotype (S11E-H Fig). We also measured transcript of the α–glucosidase *tobi*, which regulates glycogen levels and is a target of both DILPs and AKH [53], and found no effect of overexpression (not shown) or loss of function of *dilp1* at either stage (S11 I, J Fig).

### Effects of *dilp1* manipulations on metabolism in newly eclosed and young flies

To investigate whether energy reallocation during pupal development affects adult physiology and metabolism, we monitored the levels of triacylglycerids (TAG), glycogen and glucose in recently emerged and three day old *dilp* mutant and *dilp1*-overexpressing female flies (Fig 4). In newborn *dilp1* mutant flies glycogen was significantly lowered, whereas glucose and glycogen was diminished in *dilp2* mutants, while in the *dilp1/dilp2* double mutants all three compounds were decreased (Fig 4A-C). In the three-day-old flies *dilp1* and double mutants displayed reduced glycogen, whereas in *dilp1/dilp2* double mutants TAG was increased (Fig 4D-F). Using *ppl*-Ga4 to express *dilp1* we found that the only effect was a reduction of glycogen in newborn flies; at 3 or 7 days of age no effect was noted (Fig 4G-I). Thus, it appears that intact *dilp1* signaling is required for mobilization of glycogen stores in newly emerged and young flies. This supports that *dilp1* signaling in the late pupa affects metabolism and that this is carried over into the young adult.

**Fig. 4.**
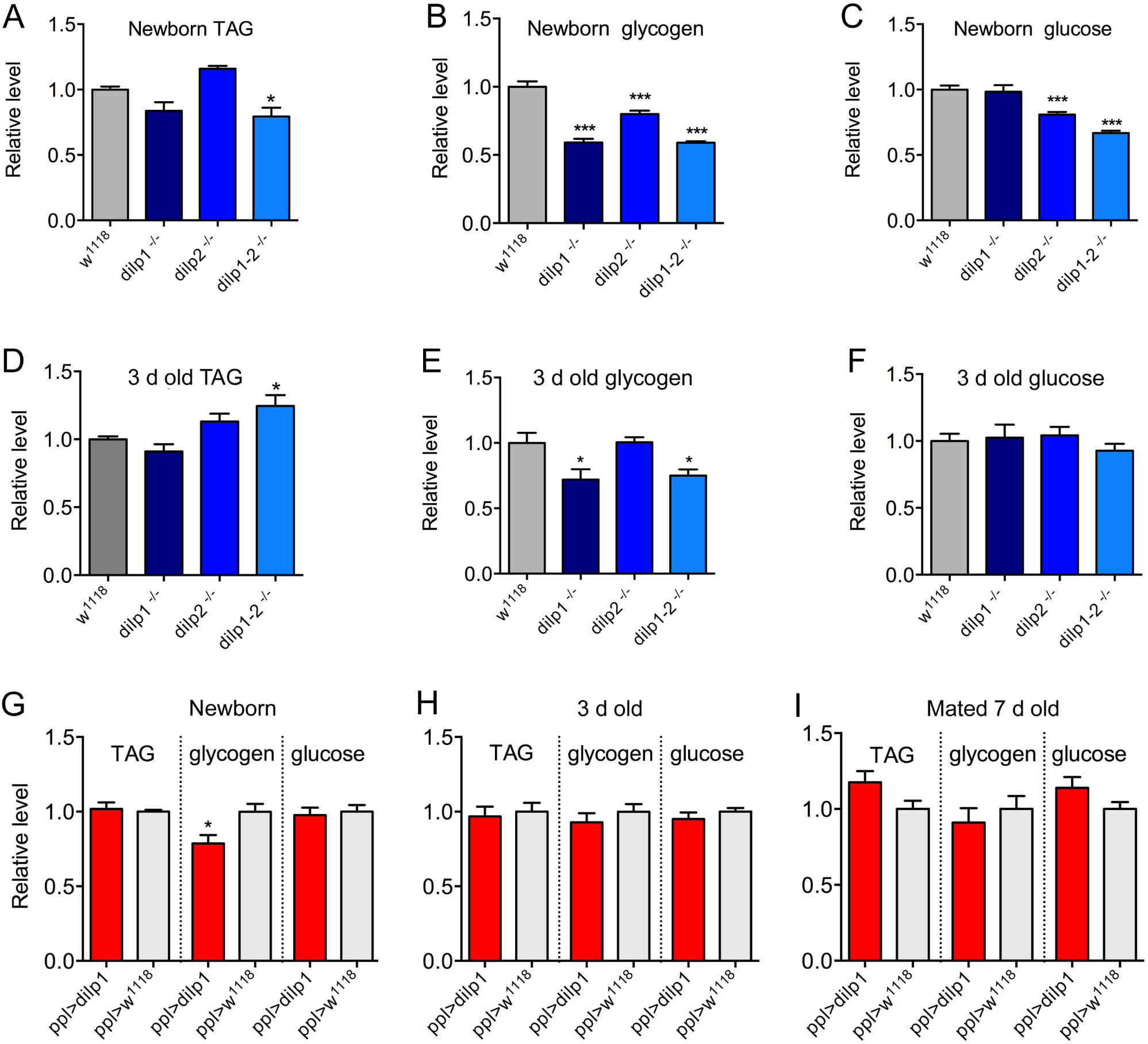
Contents in female flies of TAG, glycogen and glucose in mutants and after ectopic *dilp1* expression. **A-C.** Contents of TAG and carbohydrates in newborn mutants and controls. Note that for *dilp1* mutants only glycogen was diminished, whereas for *dilp1-2* mutants all three compounds were decreased. 8 replicates per genotype with 5-6 flies in each replicate (*p < 0.05, ***p < 0.001, one-way ANOVA followed by Tukey’s test). **D-F.** In 3 d old flies glycogen was also reduced in *dilp1* mutants and double mutants. 8 replicates per genotype with 5-6 flies in each replicate (*p < 0.05, ***p < 0.001, one-way ANOVA followed by Tukey’s test). **G-I.** Overexpression of *dilp1* in fat body (*ppl-Gal4*) only affected glycogen levels in newly hatched flies. 6-8 replicates per genotype with 5-6 flies in each. Data are presented as means ± S.E.M, (*p < 0.05, ***p < 0.001, compared to *w*^*1118*^ flies, as assessed by unpaired Students’ t-test).

### Effects of *dilp1* on adult physiology

Genetic ablation of the IPCs, which produce DILP1, 2, 3 and 5, results in enhanced starvation resistance in adult flies [21]. Thus, we asked whether the alterations of *dilp1* expression during pupal development have effects on adult physiology such as survival during starvation or desiccation (as a proxy for effects on metabolism). We investigated the starvation resistance in newly emerged, three days old and one-week-old female *dilp1, dilp2* and *dilp1/dilp2* double mutant flies. The newly eclosed *dilp1* mutant flies display strongly reduced survival during starvation and double mutants increased survival compared to control flies, whereas the starvation resistance of *dilp2* mutants is similar to the controls (Fig. 5A, Table 1). In three days old virgin flies the *dilp1* and *dilp1/dilp2* mutants display reduced survival during starvation, whereas the *dilp2* mutants perform similar to the controls (Fig 5B, Table 1). In a separate study [34] it was shown that 6-7 day old female flies display a similar response to starvation: the *dilp1/dipl2* mutants exhibit the strongest reduction in survival, followed by *dilp1* mutants that also are much less stress tolerant, whereas *dilp2* mutants and control flies perform very similar (see Table 1). Here we tested also 6-7 day old male flies and found that they survived starvation in a manner different from females with *dilp2* and double mutants displaying diminished stress resistance whereas *dilp1* mutants survive similar to controls (S12A Fig).

**Fig. 5.**
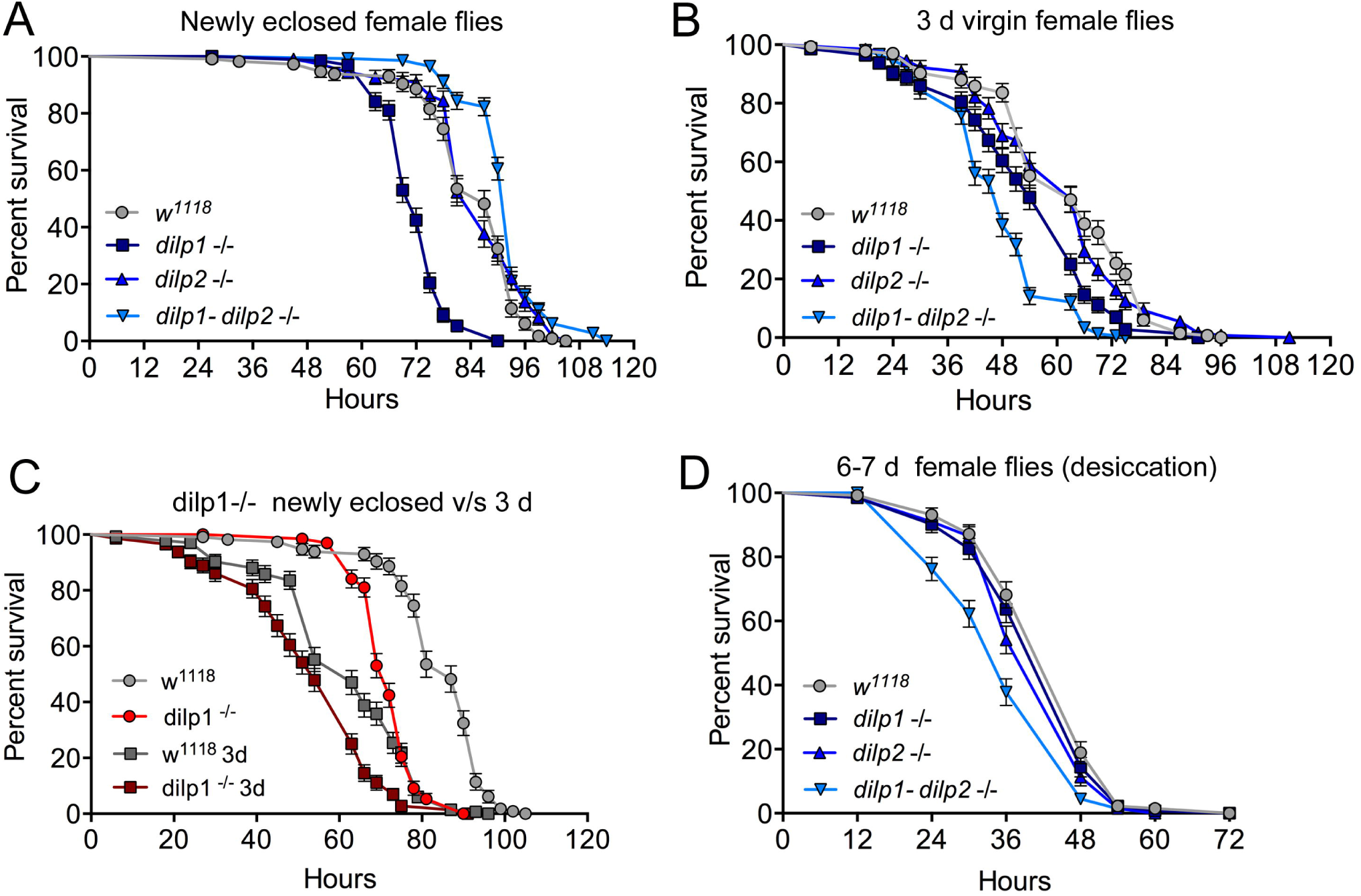
Effects of mutated *dilp* genes on adult responses to starvation and desiccation change in early adult life. **A**. In newly eclosed female flies *dilp1* mutant flies display reduced survival during starvation (p<0.001) compared to the other mutants and control. The double mutant is significantly more resistant (p<0.001). n = 109-147 flies for each genotype from three independent replicates. **B**. In three-day-old virgin female flies *dilp1-dilp2* double mutants are the least starvation resistant (p<0.001) followed by the *dilp1* mutants; n = 129-148 flies for each genotype from three independent replicates. **C.** Comparison between newly eclosed and 3 d flies exposed to starvation. Both mutants and controls survive longer as newborn flies and mutants perform worse than controls at each time point (p<0.001). n = 114-144 flies from three independent replicates. **D.** When exposed to desiccation 6-7 d old female double mutants are less resistant than the other genotypes (p<0.001), n = 132-135 flies from three independent replicates. Data are presented in survival curves and the error bars show S.E.M, as assessed by log-rank (Mantel–Cox) test].

As seen above, our data suggest a change in the response to loss of *dilp* function over the first week of adult life. It is known that newly hatched wild type flies are more resistant to starvation than slightly older flies [54]. Thus, we compared the survival during starvation in recently emerged and three day old virgin flies. As seen in Fig 5C (based on data in Fig 5A and B), recently hatched control flies (*w*^*1118*^) indeed exhibit increased starvation resistance compared to controls that were tested when three days old. Also the *dilp1* mutant flies are more starvation resistant when tested as newly hatched than as older flies, and the mutants perform less well than controls at both ages (Fig 5D). However, the most drastic change within the first week is that *dilp1* mutants yield the strongest phenotype as newborn flies and then in 3d and 6-7 d old flies the *dilp1/dilp2* mutants are the ones with the lowest stress resistance. Thus, a change in the role of *dilp1* seems to occur as the fly matures during the first few days of adult life. To provide additional evidence that *dilp1* impairs starvation resistance we performed *dilp1*-RNAi using a *dilp2*-Gal4 driver. The efficiency of the *dilp2*>*dilp1*-RNAi was tested by qPCR (S13A Fig) where a strong decrease in *dilp1*, but not *dilp2* or *dilp6* was seen. The *dilp1*-RNAi resulted in newly eclosed flies that displayed reduced survival during starvation (S13B Fig), similar to *dilp1* mutant flies.

It is also interesting to note that the diminished starvation resistance in *dilp1* and *dilp1/dilp2* mutants is opposite to the phenotype seen after IPC ablation, mutation of *dilp1-4*, or diminishing IIS by other genetic interventions [10,21,55,56]. Thus, in recently hatched flies *dilp1* appears to promote starvation resistance rather than diminishing it. Furthermore, the decreased survival during starvation in female *dilp1* mutants is the opposite of that shown in *dilp6* mutants [15], indicating that *dilp1* action is different from the other insulin-like peptides.

Next we investigated the effect of the mutations on the flies’ response to desiccation (dry starvation). One-week-old flies were put in empty vials and survival recorded. Female *dilp1/dilp2* mutants were more sensitive to desiccation than controls and the single mutants (Fig 5D). In males the double mutants also displayed higher mortality during desiccation, whereas the two single mutants were more resistant than controls (S12B Fig). Thus, there is a sex dimorphism in how the different mutants respond to both desiccation and starvation.

When overexpressing *dilp1* with the fat body driver *ppl*-Gal4 newly eclosed and 6-7 d old female flies become less resistant to starvation compared to parental controls (Fig 6A, B). However, in 6-7-day-old male flies there is no difference between controls and flies with ectopic *dilp1*, using *ppl*-and *c929*-Gal4 drivers (S13C-D Fig). We furthermore investigated starvation resistance in flies overexpressing *dilp1* in IPCs (*dilp2>dilp1*) and in most neuroendocrine cells (*c929>dilp1*) and found that in newborn flies overexpression reduced survival (Fig 6C, E), whereas in a week old flies all genotypes displayed the same survival (Fig 6D, F). Thus, in females it appears as if both knockout and over expression of *dilp1* reduces starvation resistance, maybe due to offsetting a narrow window of homeostasis. It was shown earlier that conditional knockdown of *dilp6* by RNAi during the pupal stage resulted in newborn flies with *increased* survival during starvation [15], suggesting that the effect the *dilp1* null mutation is different.

**Fig. 6.**
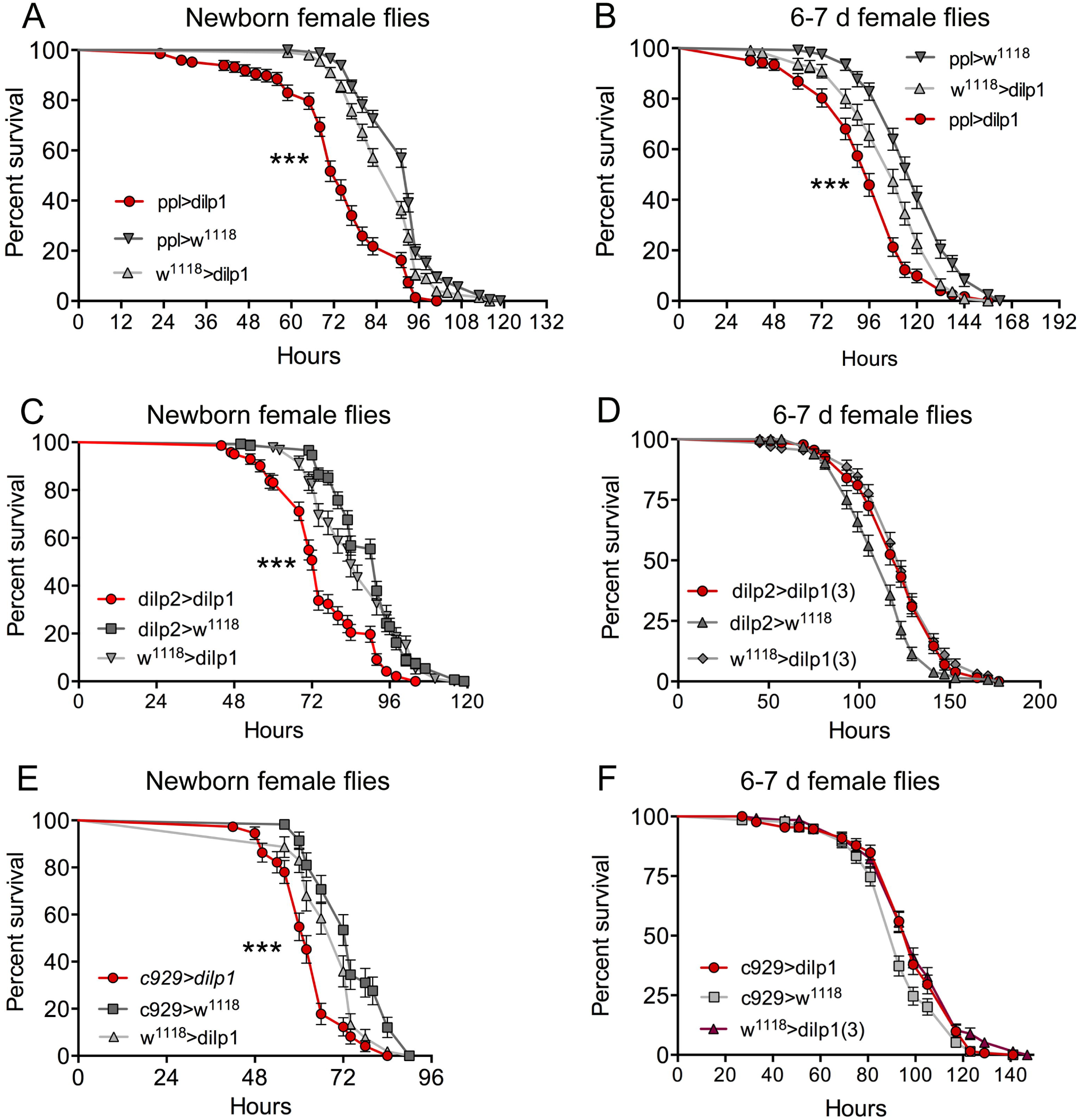
Over expression of dilp1 in the fat body affects starvation resistance in adult flies. **A** and **B.** In recently eclosed (A) and 6-7 d old (B) female flies overexpression of *dilp1* (with ppl-Gal4) leads to a decrease in survival during starvation n = 147-201 flies per genotype from three independent replicates. [***p < 0.001, as assessed by log-rank (Mantel–Cox) test]. **C** and **D.** Expressing *dilp1* in IPCs with a *dilp2*-Gal4 driver also diminishes starvation survival in newborn flies n = 92-148 flies from three independent replicates. [***p < 0.001, as assessed by log-rank (Mantel–Cox) test], but not in 6-7 d flies (n = 122-132 flies from three independent replicates). **E** and **F**. Using *c929* to drive *dilp1* in newborn and 6-7 d adult flies altered starvation resistance only in the newborn ones [***p < 0.001 as assessed by log-rank (Mantel–Cox) test, n = 132-135 flies per genotype from three independent replicates.

After ectopic expression of *dilp1* in the fat body there was an increase in food intake (cumulative data) in one-week-old flies over four days (Fig 7A), suggesting that metabolism is still altered in older flies. Since the effect of *dilp1* manipulations seems stronger in female flies we asked whether fecundity is affected by overexpression of *dilp1*. An earlier study showed that *dilp1* mutant flies are not deficient in number of eggs laid, or the viability of offspring (egg to pupal viability), although the *dilp1/dilp2* double mutants displayed a reduction in viability of these eggs [34]. Here, we expressed *dilp1* in fat body (*ppl*-Gal4) and detected an increase in number of eggs laid over 24 h in 6-7 d old flies (Fig. 7B). Both *ppl*-Gal4-and *c929*-Gal4-driven *dlip1* decreased the viability of eggs laid as monitored by numbers of eggs that developed into pupae (Fig 7C). As a comparison we noted no difference in number of eggs in 3-day-old *dilp1* mutant flies (Fig. 7D).

**Fig. 7.**
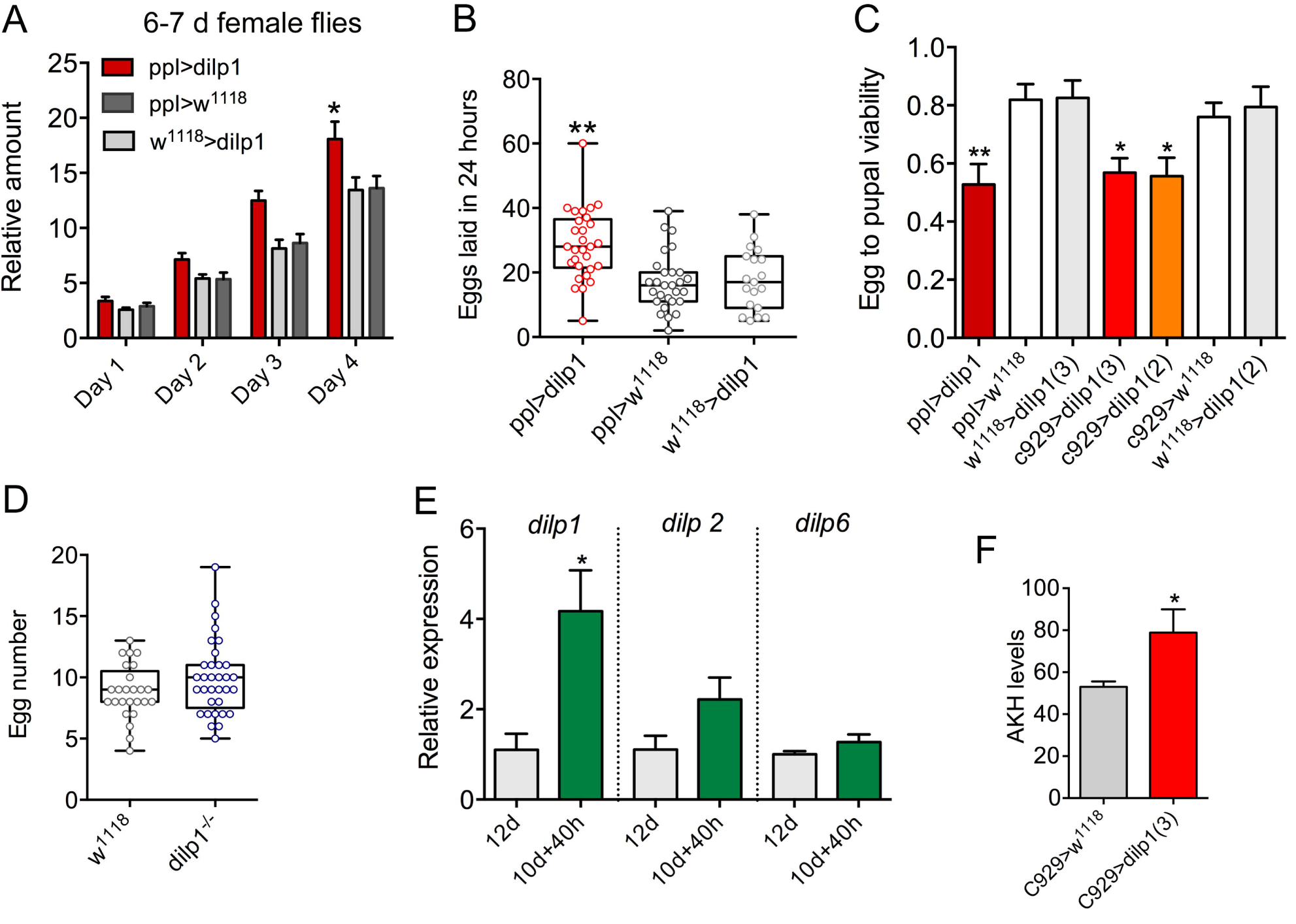
Dilp1 overexpression affects food intake and fecundity. **A.** In CAFE assay the *dilp1* overexpressing flies (6-7 d old females) display increased food intake over 4 days (cumulative data shown), Data are presented as means ± S.E.M, n = 23–24 flies from three independent replicates (*p < 0.05, two-way ANOVA followed by Tukey’s test). **B.** Number of eggs laid in 24 hours by 6-7 day old flies. We analyzed 19-29 pairs of flies from 3 replicates, (**p < 0.01, one-way ANOVA followed by Tukey’s test). **C.** The egg to pupal viability is diminished in flies with *dilp1* expressed in fat body (*ppl*-Gal4) and neuroendocrine cells [*c929*-Gal4, using two different UAS-*dilp1* (2 and 3)]. Data are presented as means ± S.E.M, more than 276 eggs from 6 replicates were monitored (*p < 0.05, unpaired Students’ t-test). **D.** Number of eggs in ovaries of 3 days old flies is not affected in *dilp1* mutants. 25-33 flies from 3 replicates were analyzed. **E.** *dilp1* mRNA is upregulated during starvation for 40 h in 10 d old adult *w*^*1118*^ flies, compared to 12 d old flies fed normal food, as monitored by qPCR. No effect was seen on *dilp2* and *dilp6* levels. Data are presented as means ± S.E.M, 3 replicates with 10 flies in each replicates were monitored (*p < 0.05, unpaired Students’ t-test). **F.** The level of adipokinetic hormone (AKH) immunolabeling in 4-5 d old female flies increased after overexpression of *dilp1* by c929>*dilp1*. 7-10 flies from 3 replicates, (*p < 0.05, unpaired Students’ t-test).

We next asked whether there is any physiological trigger of increased *dilp1* expression in adult flies, except for diapause [16] and experimental ones such as ectopic expression of sNPF or knockdown of *dilp6, dilp2* and *dilp2,3,5* [16,34,57]. Although diminished protein diet in larvae had no effect on *dilp1* expression measured by immunolabeling (not shown), we found that 40 h starvation of 10 d old flies (*w*^*1118*^) leads to a significant increase in *dilp1*, but not in *dilp2* or *dilp6* (Fig 7E). Thus, at a time (12 d) when *dilp1* is very low under normal conditions, it is upregulated four times during starvation, further suggesting that the peptide indeed plays a role also in older adult flies.

The functional homolog of glucagon in flies, AKH, plays important roles both in lipid mobilization, metabolism and regulation of lifespan [31,51,58,59]. A previous paper showed that in *dilp1* mutant flies levels of AKH were not affected [34]. Here we found that *dilp1* overexpression with the *c929*-Gal4 driver induced an increase in AKH immunolabeling in one-week-old flies (Fig 7F). Thus in adult flies (in contrast to larvae) there appears to be an interaction between *dilp1* and AKH that may underlie some of the effects of this DILP on metabolism and stress tolerance.

## Discussion

Our study indicates a role for *dilp1* in regulation of adult tissue growth during the pupal stage, as well as a function in adult physiology, especially during the first days of adult life. The experiments herein suggest that the developmental role of *dilp1* may be to ensure nutrient utilization in the pupa to support growth of adult tissues if the larva was exposed to restricted food sources. In the adult *dilp1* is upregulated during starvation and genetic gain and loss of function of *dilp1* signaling alters the flies’ survival under starvation conditions. These novel findings combined with previous data showing high levels of *dilp1* during adult reproductive diapause [16] and its role as a pro-longevity factor during aging [34] demonstrate a wide-ranging importance of this signaling system. Not only does *dilp1* expression correlate with stages of non-feeding (or reduced feeding), these stages are also associated with lack of reproductive activity, and encompass the pupae, newly eclosed flies, and diapausing flies. Under normal conditions, the diminishing *dilp1*/DILP1 expression during the first few days of adult life may be associated with a metabolic transition (fat body remodeling; [60]) and the onset of sexual maturation.

In *Drosophila*, the final body size is determined mainly during the larval feeding stage [11,12,23,29]. However, regulation of adult body size can also occur after the cessation of the feeding stage, and this process is mediated by *dilp6* acting on adult tissue growth in the pupa in an ecdysone-dependent manner [15,38]. This is likely a mechanism to ensure growth of the adult tissues if the larva is exposed to shortage of nutrition during its feeding stage. Our findings suggest that *dilp1* is another regulator of growth during the pupal stage. We show here that *dilp1* promotes organismal growth in the non-feeding pupa at the cost of stored nutrients derived from the larval stage. This is supported by RQ-data that clearly shows a shift from mixed-energy substrate energy metabolism in control flies towards almost pure lipid catabolism at the end of pupal development in the *dilp1* overexpression flies (also seen for *dilp6*). Furthermore, TAG (but not carbohydrate) levels in *dilp1* overexpression flies were clearly decreased, which likely reflects the shift in catabolic energy substrate also seen in the R/Q using respirometry. It should be noted that insects predominantly use lipids as fuel during metamorphosis [49,50] and *dilp1* overexpression increases lipid catabolism. As a consequence large *dilp1*-overexpressing flies display increased food ingestion over the first four days as adults and an altered response to starvation. Conversely *dilp1* mutants hatched as flies with significantly smaller weight. Both alterations in *dilp1* expression influence the metabolic balance in early adults as manifested in reduced starvation resistance at this stage. Our study hence suggests that *dilp1* parallels *dilp6* [15,38] in balancing adult tissue growth and storage of nutrient resources during pupal development, and thereby probably affecting adult physiology. This is interesting since *dilp6* is an IGF-like peptide that is produced in the nutrient sensing fat body [15,38], whereas the source of the insulin-like *dilp1* is the brain IPCs.

We showed earlier that young adult *dilp1* mutant flies display increased *dilp6* and vice versa [16], suggesting feedback between these two peptide hormones. This feedback appears less prominent in *dilp1* mutants during the pupal stage with no effects on *dilp2, dilp3* or *dilp6* levels. However, *dilp1* is slightly upregulated in *dilp6* mutant pupae. Furthermore, overexpression of *dilp1* in fat body or IPCs has no effect on pupal levels of *dilp2* and *dilp6*. Thus, at present we cannot postulate any compensatory changes in other DILPs in pupae with *dilp1* manipulations. However, under normal conditions *dilp6* levels are far higher than those of *dilp1* [38] (see also <u>modENCODE_mRNA-Seq_tissues</u> [61]), which could buffer the effects of changes in *dilp1* signaling.

Ectopic overexpression of *dilp1* in neuroendocrine cells or fat body not only increases growth of wings and thorax, but also increases the size of the brain and the cell bodies of several kinds of neuroendocrine cells in adult flies. However, there was no change in the size of neuronal cell bodies or CNS during larval development after overexpression of *dilp1*. Thus, taken together, our findings suggest that *dilp1*/DILP1 promotes growth mainly during the non-feeding pupal stage. Interestingly, restricted protein diet during the later larval stage diminished the body weight of adult flies more in *dilp1* mutants than in controls, similar to findings for *dilp6* [15]. This suggests that *dilp1* function is accessory to *dilp6* in maintaining growth of adult tissues in situations where larvae obtain insufficient protein in their diet.

DILPs and IIS are involved in modulating responses to starvation, desiccation and oxidative stress in *Drosophila* [see [10,21,62]]. Flies with ablated IPCs or genetically diminished IIS display increased resistance to several forms of stress, including starvation [10,21]. Conversely, overexpression of *dilp2* increases mortality in *Drosophila* [24]. We found that *dilp1* mutant flies displayed diminished starvation resistance. Both in newborn and 3 day old flies, mutation of *dilp1* decreased survival during starvation (but not in 6-7 day old ones). Curiously, overexpression of *dilp1* in the fat body also resulted in decreased survival during starvation in young and older flies. The effects on adult physiology of *dilp1* manipulations may be a consequence of the altered adult tissue growth during pupal development and associated increase in utilization of nutrient stores. Action of *dilp1* in the adult fly is also linked to reproductive diapause in females, where feeding is strongly reduced [63], and both peptide and transcript are upregulated [16]. Related to this we found here that *dilp1* mRNA is upregulated during starvation in 12 d old flies. Furthermore, it was shown that expression of *dilp1* increases lifespan in *dilp1-dilp2* double mutants, suggesting that loss of *dilp2* induces *dilp1* as a factor that promotes longevity [34]. Thus, *dilp1* activity is beneficial also during adult life, even though its expression under normal conditions is very low [15,16,38]. This pro-longevity effect of *dilp1* is in contrast to *dilp2, 3 and 5* and the mechanisms behind this effect are of great interest to unveil.

A previous study showed that in wild-type (Canton S) *Drosophila* DILP1 expression in young adults is sex-dimorphic with higher levels in females [16]. In line with this, we show here that increase in body weight the first week of adult life occurs only in female *dilp1* mutant flies, and also that starvation survival in one-week-old flies is diminished only in females. Finally, we found that *dilp1* over expression specifically decreased starvation resistance only in female flies. Thus, taken together, we found earlier that *dilp1* displays a sex-specific expression [16] and here we show sex-specific function in young adult *Drosophila*, and the *dilp1* mutation affects body weight of newly eclosed flies mainly in females. It is tempting to speculate that the more prominent role of *dilp1* in female flies is linked to metabolism associated with reproductive physiology and early ovary maturation, which is also reflected in the *dilp1* upregulation during reproductive diapause [16]. In fact, we show here that egg-laying increased after *dilp1* overepression, and an earlier study demonstrated a decreased egg laying in dilp1 mutant flies [16].

This study demonstrates that *dilp1* promotes growth of adult tissues during the pupal stage, and in females it influences starvation resistance during the young adult stage, and affects fecundity. Like *dilp6*, perhaps *dilp1* acts as a signal promoted by nutrient shortage during the late larval stage to ensure growth of adult tissues by recruiting nutrient stores from larval fat body. This in turn results in depleted pupal-derived nutrient stores in young adults. Thus, IPC-derived *dilp1* displays several similarities to the fat body-produced *dilp6*, including temporal expression pattern, growth promotion, effects on adult stress resistance and lifespan. Additionally *dilp1* may play a role in regulation of nutrient utilization/metabolism during the first few days of adult life, especially in females. At this time larval fat body is still present and utilized as energy fuel/nutrient store [54] and also contribute to egg development [64]. Curiously, there is a change in the action of DILP1 between the pupal and adult stages from being a stimulator of growth (agonist of dInR) in pupae, to acting opposite to DILP2 and other DILPs in adults in regulation of lifespan and stress responses. It is not known what mechanism is behind this switch in function of DILP1 signaling, but one possibility is that DILP1 acts via different signaling pathways downstream the dInR in pupae and adults. One obvious difference between these two stages is the presence of larval fat body in the pupa and first few days of adults and its replacement by functional adult fat body in later stages [37,54]. Also there seems to be a difference in the interactions with AKH signaling. During pupal development we did not see any effect of *dilp1* on transcripts of *Akh* or *tobi*, whereas in adult flies *Akh* expression is induced by *dilp1* [34]. This is in agreement with earlier work, which showed that AKH plays no role in development or lipid and carbohydrate metabolism in the pupa [51]. In the future it would be interesting to investigate whether DILP1 acts differently on larval/pupal and adult fat body, or act on different downstream signaling in the two stages of the life cycle, and whether *dilp1* and *dilp6* interact to regulate growth and metabolism in *Drosophila*.

## Experimental procedures

### Fly lines and husbandry

Parental flies were reared and maintained at 18°C with 12:12 Light:Dark cycle on food based on a recipe from Bloomington *Drosophila* Stock Center (BDSC) (http://fly-stocks.bio.indiana.edu/Fly_Work/media-recipes/bloomfood.htm). The experimental flies were reared and maintained at 25°C, with 12:12 Light:Dark cycle on an agar-based diet with 10% sugar and 5% dry yeast.

The following Gal4 lines were used in this study: *dilp2*-Gal4 [[18] from E. Rulifson, Stanford, CA], *Pdf*-Gal4 (obtained from BDSC, Bloomington, IN), *ppl*-Gal4 [[65] from M.J. Pankratz, Bonn, Germany], *To*-Gal4 [[66] from B. Dauwalder, Houston, TX], *c929*-Gal4 [[46] from Paul H. Taghert], yw; UAS-*dilp6*, and yw; UAS-*dilp2;+* [[23] from H. Stocker, Zürich, Switzerland]. Several UAS-*dilp1* lines were produced for a previous study [34] and two of them, UAS-*dilp1* (II) and UAS-*dilp1* (III), were used here. UAS-*dilp1*-RNAi flies were from Vienna *Drosophila* Resource Center (VDRC), Vienna, Austria. As controls we used *w*^*1118*^ or *yw* obtained from BDSC, crossed to Gal4 and UAS lines. All flies (except yw; UAS-*dilp6*, and yw; UAS-*dilp2;+*) were backcrossed to *w*^*1118*^ for at least 6 generations.

We used a double null mutation of *dilp1/dilp2* that was previously generated by homologous recombination and verified as described by Post et al. [34]. Also single *dilp1* and *dilp2* null mutants were employed. We refer to these three null mutants as *dilp1, dilp2* and *dilp1/dilp2* mutants for simplicity. As described earlier [34], these were obtained from BDSC and a residual w+ marker was Cre excised followed by chromosomal exchange to remove *yw* markers on chromosomes 2 and X.

To generate a recombinant *dilp6;;dilp1* double mutant, the *dilp1* and *dilp6*^*68*^ mutants [10] were used for crossing with a double balancer fly, 4E10D/FM7,dfd;;Vno/TM3,dfd, obtained from Dr. Vasilios Tsarouhas (Stockholm University). The efficiency of the *dilp6;;dilp1* double mutant was validated by qPCR.

### Antisera and immunocytochemistry

For immunolabeling, tissues from larvae or female adults were dissected in chilled 0.1 M phosphate buffered saline (PBS). They were then fixed for 4 hours in ice-cold 4% paraformaldehyde (PFA) in PBS, and subsequently rinsed in PBS three times for 1 h. Incubation with primary antiserum was performed for 48 h at 4°C with gentle agitation. After rinse in PBS with 0.25% Triton-X 100 (PBS-Tx) four times, the tissues were incubated with secondary antibody for 48 h at 4°C. After a thorough wash in PBS-Tx, tissues were mounted in 80% glycerol with 0.1 M PBS.

The following primary antisera were used: Rabbit or guinea pig antiserum to part of the C-peptide of DILP1 diluted 1:10000 [16]. Rabbit antisera to A-chains of DILP2 and DILP3 [67] and part of the C-peptide of DILP5 [68] all at a dilution of 1:2000, rabbit anti-AKH (1:1000) from M.R. Brown, Athens, GA, rabbit anti-pigment-dispersing hormone (1:3000) from H. Dircksen, Stockholm, Sweden [69], rabbit antiserum to cockroach leucokinin I (LK I) at 1:2000 [70], mouse anti-green fluorescent protein (GFP) at 1:000 (RRID: AB_221568, Invitrogen, Carlsbad, CA).

The following secondary antisera were used: goat anti-rabbit Alexa 546, goat anti-rabbit Alexa 488, and goat anti-mouse Alexa 488 (all from Invitrogen). Cy3-tagged goat anti-guinea pig antiserum (Jackson ImmunoResearch, West Grove, PA). All were used at a dilution of 1:1000.

### Image analysis

Images were captured with a Zeiss LSM 780 confocal microscope (Jena, Germany) using 10×, 20× and 40× oil immersion objectives. The projections of z-stacks were processed using Fiji (https://imagej.nih.gov/ij/). The cell body outlines were extracted manually and the size and staining intensity were determined using ImageJ (https://imagej.nih.gov/ij/). The background intensity for all samples was recorded by randomly selecting three small regions near the cell body of interest. The final intensity value of the cell bodies was determined by subtracting the background intensity.

Images of pupae, adult flies and fly wings were captured with a Leica EZ4HD light microscope (Wetzlar, Germany). The size of the adult fly body and wings were determined using Fiji. The pupal volume (v) was calculated using the equation v = 4/3 π (L/2) × (l/2)^2^, in which L = length and l = width [71]. Thorax length was measured from the posterior tip of the scutellum to the base of the most anterior point of the humeral bristle.

### Pupariation time, egg to pupae viability and adult body weight

To determine time to pupariation, 6-7 day old adult females were crossed in the evening. The following morning, adult flies were transferred to vials with fresh food on which they were allowed to lay eggs for four hours. Two hours after the initiation of egg laying was considered time “0”, and thereafter the number of pupae was monitored at 6 or 12-hour intervals. To investigate the viability of egg to pupae formation, one pair of 6-7 day old adult flies was allowed to lay eggs for 24 hours after which the total number of eggs was counted. Subsequently, the total number of pupae was counted and the viability of egg to pupae was determined as pupa number/egg number × 100%. The body weight (wet weight) of single adult flies was determined using a Mettler Toledo MT5 microbalance (Columbus, USA). The number of eggs of stage 10-14 in ovaries was counted in 3-day-old flies.

### Starvation survival assay

Newly hatched and mated 6-7 day old adults were used for starvation resistance experiments. For newly hatched flies, we collected virgin flies every 4 hours, to be used for starvation experiments. The flies were kept in vials containing 5 ml of 0.5% aqueous agarose (A2929, Sigma-Aldrich). The number of dead flies was counted at least every 12 hours until all the flies were dead. At least 110 flies from 3 replicates were used for the analysis.

### Capillary feeding (CAFE) assay

Food intake was measured using a slightly modified capillary feeding (CAFE) assay following Ja et al. [72]. In brief, female flies were placed into 1.5-ml Eppendorf micro centrifuge tubes with an inserted capillary tube (5 µl, Sigma) containing 5% sucrose, 2% yeast extract and 0.1% propionic acid. To estimate evaporation, three food-filled capillaries were inserted in identical tubes without flies. The final food intake was determined by calculating the decrease in food level minus the average decrease in the three control capillaries. Food consumption was measured daily and calculated cumulatively over four consecutive days. For this assay we used 8-10 flies in each of three biological replicates.

### Quantitative real-time PCR (qPCR)

Total RNA was extracted from whole bodies of middle or late pupal stages of *Drosophila* by using Trizol-chloroform (Sigma-Aldrich). Quality and concentration of the RNA were determined with a NanoDrop 2000 spectrophotometer (Thermo Scientific). The concentration of the RNA was adjusted to 400 ng/µl. Totally 2ug RNA was used for cDNA synthesis. The cDNA syntheses were performed by using random hexamer primer (Thermo Scientific) and RevertAid reverse transcriptase (Thermo Scientific). The cDNA products were then diluted 10 times and applied for qPCR using a StepOnePlus™ instrument (Applied Biosystem, USA) and SensiFAST SYBR Hi-ROX Kit (Bioline) followed the protocol from the manufacturer. The mRNA abundance was normalized to ribosomal protein (rp49) levels in the same samples. Relative expression values were determined by the 2^-ΔΔCT^ method [73]. The sequences of primers used for qPCR were those used previously [16,34,74]:

dilp1 F: CGGAAACCACAAACTCTGCG

dilp1 R :CCCAGCAAGCTTTCACGTTT

dilp2 F: AGCAAGCCTTTGTCCTTCATCTC

dilp2 R: ACACCATACTCAGCACCTCGTTG;

dilp6 F: CCCTTGGCGATGTATTTCCCAACA

dilp6 R: CCGACTTGCAGCACAAATCGGTTA

akh F: GCGAAGTCCTCATTGCAGCCGT

akh R: CCAATCCGGCGAGAAGGTCAATTGA

*tobi* F: CCACCAAGCGAGACATTTACC

*tobi* R: GAGCGGCGTAGTCCATCAC

*bmm* F: GGT CCC TTC AGT CCC TCC TT

*bmm* R: GCT TGT GAG CAT CGT CTG GT

rp49 F: ATCGGTTACGGATCGAACAA

rp49 R: GACAATCTCCTTGCGCTTCT

### Metabolite quantification

Glycogen and triacyl glyceride (TAG) levels were assayed as previously described [34,75,76]. For glycogen assays, 5-6 adult female flies per sample were homogenized in PBS and quantified using the Infinity Glucose Hexokinase reagent by spectrophotometry. For TAG assays, 5-6 adult female flies per sample were homogenized in PBS + 0.05% TBS-T and quantified using the Infinity Triglycerides reagent by spectrophotometry. The fly lysate protein levels were determined by BCA assay (Thermo Fisher) and metabolite levels were normalized to protein level.

To measure the amount of TAG during late pupal stages, 6 replicates with 4 pupae in each were collected and then homogenized in PBS + 0.05% Triton-X 100 with a tissuelyser II from Qiagen. The TAG levels was determined with a Liquick Cor-TG diagnostic kit (Cormay, Poland) using a linear regression coefficient from a standard curve made with 2.2 µg/µl TAG standard (Cormay, Poland). Absorbance of samples was measured at 550 nm with a micro-plate reader (Thermo scientific). Data are expressed as micrograms of TAG related to protein levels. Protein levels were determined using a Bradford assay according to Diop et al. [77].

### Dynamic injection respirometry

Carbon dioxide (CO_2_) production and oxygen (O_2_) consumption of individual pupae of both sexes were measured during pupal development at 25°C to assess metabolic rate (MR) as described previously [49]. Pupae were placed in 1 ml syringes (i.e. respirometry chambers) that were filled with air scrubbed of CO_2_ with ascarite (Acros Organics, USA) that then passed through filtered acidified water (pH < 4.5, checked weekly), closed with three-way luer valves, and kept for roughly 24 hours at 25°C with 12:12 Light:Dark cycle. An empty syringe served as control. CO_2_ production was measured using a Sable Systems (Las Vegas, NV, USA) differential respirometry setup. Two independent lines of outdoor air scrubbed of H_2_O and CO_2_, using drierite (WA Hammond Drierite, USA) and ascarite scrubbers respectively, were pushed at a steady rate of 150 ml min^-1^ using a SS-4 pump (Sable Systems) and two separate mass flow controllers (840 Series; Sierra Instruments Inc, California, USA). The syringes containing pupae were placed after the mass valve controllers in the first line (sample) and 0.45 ml pushed into the airflow. The push rate was recorded through a second flowmeter downstream of the syringe and approximated a flow rate of 162 ml min^-1^ downstream of the syringe. The line was then scrubbed of H_2_O with drierite and entered the sample line of a Li-7000 CO_2_ analyser (LiCor, Lincoln, NE, USA). The second line (reference) proceeded the same way, mimicking the exact length of the sample line (including an empty measurement chamber), entering the reference line of the CO_2_ analyser. The lines then proceeded through a second set of ascarite CO_2_ scrubbers and entered an Oxzilla FC-2 O_2_ analyser (Sable Systems) after which air was ejected. Preliminary measurements were performed to ensure stability of flow rate through either channel by measuring the flow rate of air ejected from the O_2_ analyser. After the measurement pupae were weighed using a Mettler Toledo MT5 microbalance (Columbus, USA) and left at 25°C with 12:12 Light:Dark cycle until adult eclosion, at which point they were sexed.

Differential CO_2_ and O_2_ were calculated by subtracting the output of the reference line from the output of the sample line. For all measurements sampling rate was 1 Hz. In the program Expedata (version 1.9.10) the raw output was baseline corrected against the reference line value, fractioned and multiplied with flow rate to yield CO_2_ and O_2_ in ml min^-1^ [78]. The values were then corrected by subtracting the readings from the empty control syringe from the sample values. MR was calculated by first integrating the fractioned CO_2_ and O_2_ (ml min^-1^) values against time to yield CO_2_ and O_2_ in ml produced while pupae were in the syringes. Next *V*CO_2_ and *V*O_2_ were corrected by accounting for the fraction of air that was still left in the syringe and the time spent in the syringe using the formula (only calculation for *V*CO_2_ is shown) *V*CO_2_ = (CO_2_ × (0.6 / 0.45)) / hours in syringe (Lighton, 2008). Then the respiratory quotient (RQ) was calculated as RQ = *V*CO_2_ / *V*O_2_. RQ values provide an estimate on what energy source is being catabolized to fuel metabolism [48]. MR (in Watts = Joules s^-1^) was converted from *V*O_2_ using the formula MR = (*V*O_2_ × (16 + (5.164 × RQ))) / (60 × 60) (Lighton 2008) and finally divided by body weight in mg to yield MR mg^-1^.

In the present study we monitored single identified individuals throughout pupal development, and sexed them after eclosion. For the vast majority eclosion was successful and therefore we could use the true weight of the individual for the calculation above. However, for individuals that failed to eclose properly we instead used the average weight for that sex and treatment to calculate MR.

### Statistical analysis

All results are presented as means ± SEM. We first investigated normality of data using Shapiro-Wilk’s normality test, then used one-way analysis of variance (ANOVA) or Student’s t-test, followed by Tukey’s multiple comparisons test. Lifespan data were subjected to survival analysis (Log rank tests with Mantel-Cox post-test) and presented as survival curves.

For the respirometry data we used the natural logarithm of MR mg^-1^ due to deviations from normality. A factorial two-way ANOVA was used with MR mg^-1^ or RQ as dependent variable, and sex and treatment as factorial explanatory variables. Non-significant interactions and main effects were removed from final models [79]. The respirometry data were analyzed with the IBM SPSS statistics 23.0 (IBM SPSS Inc., Chicago, IL, USA) statistical software package. Prism GraphPad version 6.00 (La Jolla, CA, USA) was used for generating all the graphs.

## Acknowledgements

We thank the Bloomington *Drosophila* Stock Center (Bloomington, IN), the Vienna *Drosophila* Resource Center (Vienna, Austria) and Drs. M.J. Pankratz, B. Dauwalder, V. Monnier, P.H. Taghert, V. Tsarouhas and H. Stocker for flies, and Dr. M. R. Brown for AKH antiserum.

## Supplemental material figures

S1 Fig. Evaluation of mutant efficiency. **A**. qPCR reveals that in stage P8-9 pupae the *dilp1* and *dilp1/dilp2* mutants display *dilp1* levels that are close to zero, whereas in the *dilp6* mutant *dilp1* is upregulated and in *dilp2* mutant slightly reduced. **B.** In the *dilp2* and *dilp1/dilp2* mutants *dilp2* levels are not detectable. **C.** The *dilp6* levels are only affected in the *dilp6* mutants. Data are presented as means ± S.E.M, n = 6 replicates for each genotype with 6 pupae in each replicate. (*p < 0.05, compared with w^1118^ flies, unpaired Students’ t-test). **D.** Using immunocytochemistry with antisera to DILP1-3 it can be shown that labeling of IPCs in 1-week-old flies is not detectable for anti-DILP1 in *dilp1* and double mutants and for DILP2 in *dilp2* and double mutants. DILP3 is upregulated in *dilp2* mutants. **E-G**. Quantification of immunofluorescence shows that DILP1 labeling is not affected in *dilp2* mutants (E), DILP2 is increased in *dilp1* mutants (F) and DILP3 strongly increased only in *dilp2* mutants (G). Data are presented as means ± S.E.M, n = 9-12 flies from 3 replicates. (**p < 0.01, ***p < 0.001, compared with w^1118^ flies, unpaired Students’ t-test).

**S2 Fig.** Recombinant *dilp1/dilp6* mutant flies display reduced body mass. **A.** Transcripts of *dilp1* and *dilp6* in one-day-old *dilp1/dilp6* mutant flies. Data are presented as means ± S.E.M, n = 3 replicates for each genotype with 6 pupae in each replicate. (**p < 0.001, ***p < 0.0001, compared with w^1118^ flies, unpaired Students’ t-test). **B.** Body weights are significantly reduced in single mutants and recombinant double mutants, but no additive effect of the double mutation was detected. (n = 11–16 flies per genotype from three replicates, One-way ANOVA followed by Tukey’s test).

**S3 Fig.** Verification of ectopic *dilp1* expression by DILP1 immunolabeling. **A.** After *dilp2*-Gal4-driven *dilp1* expression strong DILP1 immunolabeling can be detected in IPCs of 3^rd^ instar larvae as well as 1 and 3 week old adults, but not in controls (*dilp2*>w^1118^). **B.** Quantification of DILP1 immunofluorescence in IPCs of one-week-old adults, using two different UAS-*dilp1* (2 and 3). Data are presented as means ± S.E.M, n = 5-7 flies from 3 replicates. (***p < 0.001, compared with control flies, unpaired Students’ t-test). **C.** Using the c929 driver DILP1 immunolabeling can be detected in numerous neuroendocrine cells in the CNS of larvae and brain of adults, but not in controls (c929>w^1118^). **D.** Using two different fat body Gal4 drivers (*ppl* and *to*) DILP1 immunolabeling can be detected in adipocytes.

**S4 Fig.** Verification of ectopic *dilp1* expression by qPCR in stage P8-9 pupae. **A.** Using the fat body Gal4 drivers *ppl* and *to* a drastic increase of *dilp1* transcript was seen. **B.** The *dilp2* level was diminished after ppl-driven *dilp1*. **C.** No significant effect was seen on *dilp6* levels after *dilp1* expression. **D-F.** Driving *dilp1* in IPCs with *dilp2*-Gal4 drastically increases *dilp1*, but has no effect on *dilp2* or *dilp6*. Data are presented as means ± S.E.M, n = 5-6 replicates per genotype with 10 pupae in each replicate. (*p < 0.05, **p < 0.01, ***p < 0.01, compared with w^1118^ flies, unpaired Students’ t-test).

**S5 Fig.** Effects of ectopic *dilp1* expression on peptide levels of DILPs in one-week-old adults. **A**. Expressing *dilp1* in IPCs (*dilp2>dilp1*) increases DILP2 immunolabeling and decreases DILP3. **B** and **C**. Quantification of immunolabeling. Data are presented as means ± S.E.M, n = 7-10 per genotype from 3 replicates. (**p < 0.01, compared with w^1118^ flies, unpaired Students’ t-test)**. D.** Using the broader *c929*-Gal4 to drive *dilp1* the DILP5 immunolabeling of IPCs increase. **E.** Quantification of DILP5 immunolabeling. Data are presented as means ± S.E.M, n = 9-12 from 3 replicates. (**p < 0.01, compared with w^1118^ flies, unpaired Students’ t-test).

**S6 Fig.** Effects of ectopic *dilp1* expression on body weight and organismal size. **A.** Expressing *dilp1* in the fat body (*ppl*-Gal4) of male flies leads to increased weight compared to controls in both young and older flies. However, in contrast to female flies, shown in Fig. 2K, there is no gain in weight over the first 5-6 days as adults, rather a decrease. Data are presented as medians ± range, n = 14–25 flies from three independent replicates (*p < 0.05, **p < 0.01, ***p < 0.001, two-way ANOVA followed with Tukey’s test). **B.** Driving *dilp1* in IPCs with *dilp2*-Gal4 in females increases the weight compared to controls in older flies. Data are presented as medians ± range, n = 14–23 flies from three independent replicates (*p < 0.05, **p < 0.01, two-way ANOVA followed with Tukey’s test). **C**. Driving *dilp1* in IPCs increases the weight of one-day-old and 6-7 day old male flies, compared to both controls. Furthermore the younger flies weigh more than the older ones for all genotypes. Data are presented as medians ± range, n = 14–24 flies per genotype from three independent replicates (**p < 0.01, ***p < 0.001, two-way ANOVA followed with Tukey’s test). **D** and **E.** Using *to*-Gal4 the body masses show the same patterns as with *ppl*-Gal4 (Fig. 2K and S5A Fig), where body masses increase after *dilp1* over expression, and in females there is an additional weight gain over the first 5-6 days. The following days (13-14 d) no additional increase is seen. Data are presented as medians ± range, n = 9–27 flies per genotype from three independent replicates (*p < 0.05, **p < 0.01, ***p < 0.001, two-way ANOVA followed with Tukey’s test). **F-H.** The *dilp1* expression obtained with the *to*-Gal4 does not result in a significant increase in wing area, (n = 16–22 flies per genotype from three replicates, One-way ANOVA followed with Tukey’s test), whereas thorax length increased slightly (n = 19-32 flies per genotype (*p < 0.05 unpaired Students’ t-test), but no effect on pupal volume (n = 29 flies per genotype from three replicates, One-way ANOVA followed with Tukey’s test).

**S7 Fig.** Effects of *dilp1* expression on weight, wing area and food intake. **A** The body weight increased in male flies after ectopic *dilp1* expression with *c929*-Gal4, *** p<0.001, data are presented as means ± S.E.M, n = 16–29 flies per genotype from three independent replicates (One-way ANOVA followed with Tukey’s test). **B**. The wing area is not affected by *c929*-driven *dilp1* expression. Data are presented as means ± S.E.M, n = 15 flies from three independent replicates (One-way ANOVA followed with Tukey’s test). **C.** Driving *dilp1* with *dilp2*-and *c929*-Gal4 does not affect food intake. Data are presented as means ± S.E.M, n = 24 flies from three independent replicates (two-way ANOVA followed with Tukey’s test). dent replicates, as assessed by log-rank (Mantel–Cox) test.

**S8 Fig.** The brain and neuronal cell bodies grow after *dilp1* overexpression in neuroendocrine cells. **A-C.** Using the *c929*-Gal4 line to drive *dilp1* in neuroendocrine cells leads to increased size of the cell bodies of DILP2 immunolabeled insulin producing cells (A1-A3), PDF labeled l-LNv clock neurons (B1-B3) and abdominal leucokinin (LK) immunoreactive neurons, ABLK (C1-C3). **D.** The entire brain also increases in size in *c929>dilp1* flies. **E.** Expression of *dilp1* in IPCs with the *dilp2*-Gal4 line is not sufficient to obtain an increase in size of IPCs. **F1-F3.** Expression of *dilp1* in the fat body (*ppl*-Gal4) increases the size of the l-LNv clock neurons and the entire brain (**G**). Data are presented as means ± S.E.M, n = 8–10 samples for each genotype from three independent replicates (*p < 0.05, **p < 0.01, ***p < 0.001, as assessed by unpaired Students’ t-test).

**S9 Fig.** Ectopic expression of *dilp1* in clock neurons or larval neuroendocrine cells does not affect cell size. **A-D.** Expression of *dilp1* with the clock neuron driver *pdf*-Gal4 does not affect the size of the PDF-immunolabeled large LN_v_s quantified in B. The brain size is also not affected (C). However the PDF immunolabeling is strongly increased (D). Data are presented as means ± S.E.M, n = 8 for each genotype from 3 replicates. (**p < 0.01, compared with w^1118^ flies, unpaired Students’ t-test). **E.** Ectopic expression of *dilp1* with the c929-Gal4 line does not affect the size of leucokinin (LK)-immunolabeled neuronal cell bodies in the third instar larvae (quantified in F) or the size of the larval CNS (G). Data are presented as means ± S.E.M, n = 6-9 for each genotype from 3 replicates.

**S10 Fig.** Ectopic expression of *dilp1* in glial cells with *repo*-Gal4 does not affect growth of neuronal cell bodies. **A.** DILP1 immunolabeling appears in cells after *Repo>dilp1*, but has no effect on the size of l-LNv clock neurons labeled with anti-PDF (quantified in **B**). Data are presented as means ± S.E.M, n = 9 for each genotype from 3 replicates.

**S11 Fig.** Overexpression and mutation of *dilps* have little effect on AKH signaling as determined by qPCR. **A -J.** Transcripts of AKH (*Akh*), brummer lipase (*bmm*) and the glucosidase target of brain insulin (*tobi*) were measured by qPCR in different genotypes at two stages of pupal development: two day old and 4 day old. For overexpression in fat body we used *ppl*-and *to*-Gal4 drivers. We analyzed 3 replicates with 6 pupae in each replicates (*p < 0.05, one-way ANOVA followed by Tukey’s test). **K.** Glucose levels were determined in 4 d pupae; 6 replicates per genotype with 4 pupae in each replicate (No significant differences; analysis by one-way ANOVA followed by Tukey’s test). This panel is associated with Fig. 3D – F.

**S12 Fig.** Effect on starvation and desiccation in male *dilp* mutant flies. **A.** In 6-7 days old male flies *dilp1-dilp2* mutants are least resistant to starvation (p<0.001), followed by *dilp2* mutants (p<0.001), whereas *dilp1* mutants perform as controls; n = 125-141 flies from three independent replicates. However 6-7 d female flies perform as 3 d virgin females (see [34] and Table 1). **B.** In males double mutants are less (p<0.001), and the other two mutants more resistant (p<0.001) to desiccation than controls, n = 134-135 flies from three independent replicates. Data are presented in survival curves and the error bars means S.E.M, as assessed by log-rank (Mantel–Cox) test].

**S13 Fig.** Targeted *dilp1*-RNAi in IPCs reduces survival in flies exposed to starvation. **A.** The efficiency of *dilp2*>*dilp1*-RNAi on *dilp1* levels was monitored by qPCR. A strong reduction in *dilp1* was noted, but no effect was seen on levels of *dilp2* or *dilp6*. Data are presented as means ± S.E.M, n = 3 replicates per genotype with 10 pupae in each replicate. (*p < 0.05, compared with control flies, unpaired Students’ t-test). **B.** In newly eclosed female flies *dilp2*>*dilp1*-RNAi resulted in reduced survival during starvation. n = 148-170 flies from three independent replicates. Data are presented in survival curves and the error bars means S.E.M [***p < 0.001, as assessed by log-rank (Mantel–Cox) test]. **C.** In 6-7 d old males *dilp1* overexpression in fat body (*ppl*-Gal4) has no effect on starvation response. n = 117-128 flies from three independent replicates. **E**. c929-driven *dilp1* does not affect the response to starvation, n = 132-135 flies per genotype from three independent replicates.

