## Supplementary figures and images for "*Drosophila* insulin-like peptide 1 (DILP1) promotes organismal growth and catabolic energy metabolism during the non-feeding pupal stage"

### S Fig. 1.jpg

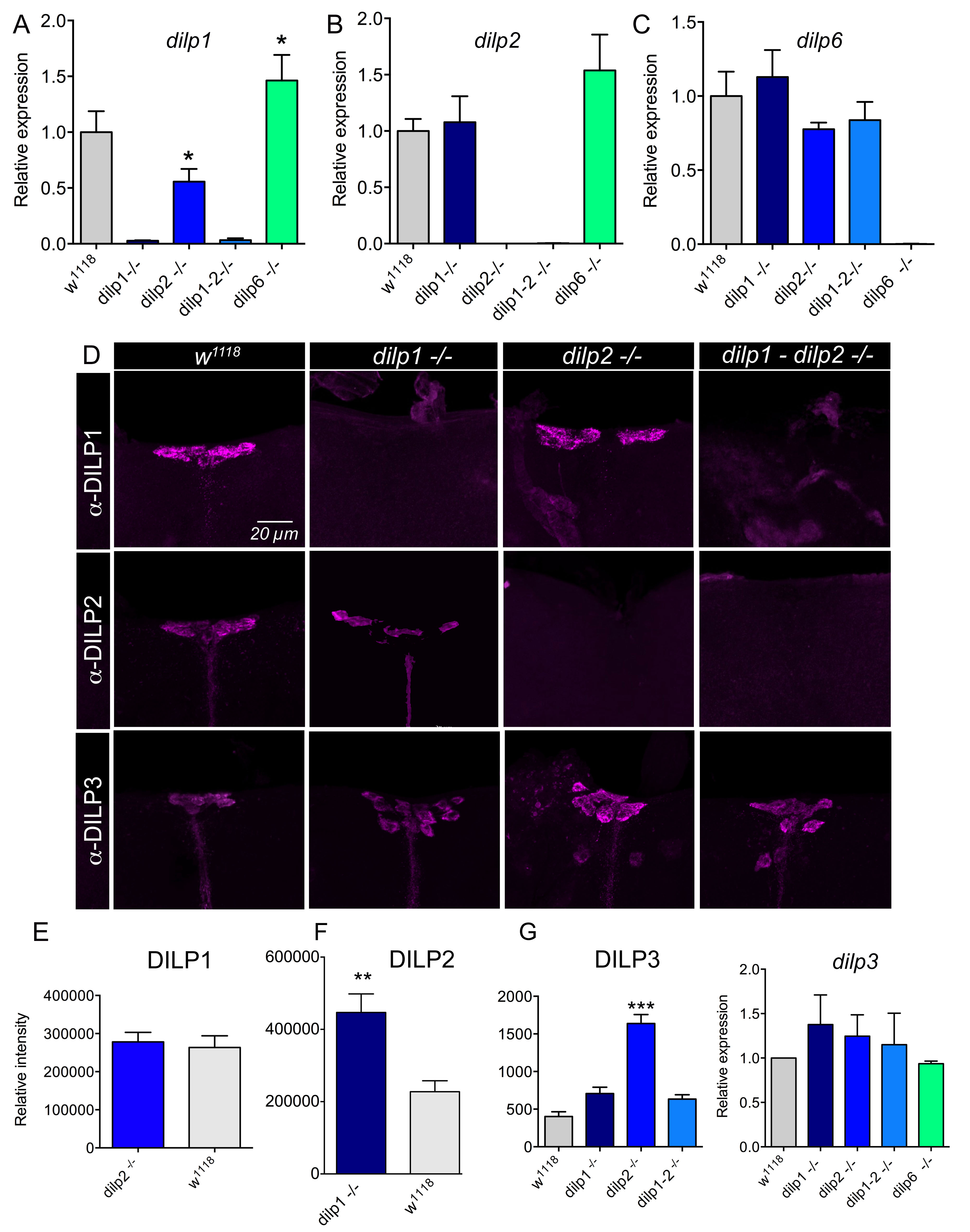

### S Fig. 2.jpg

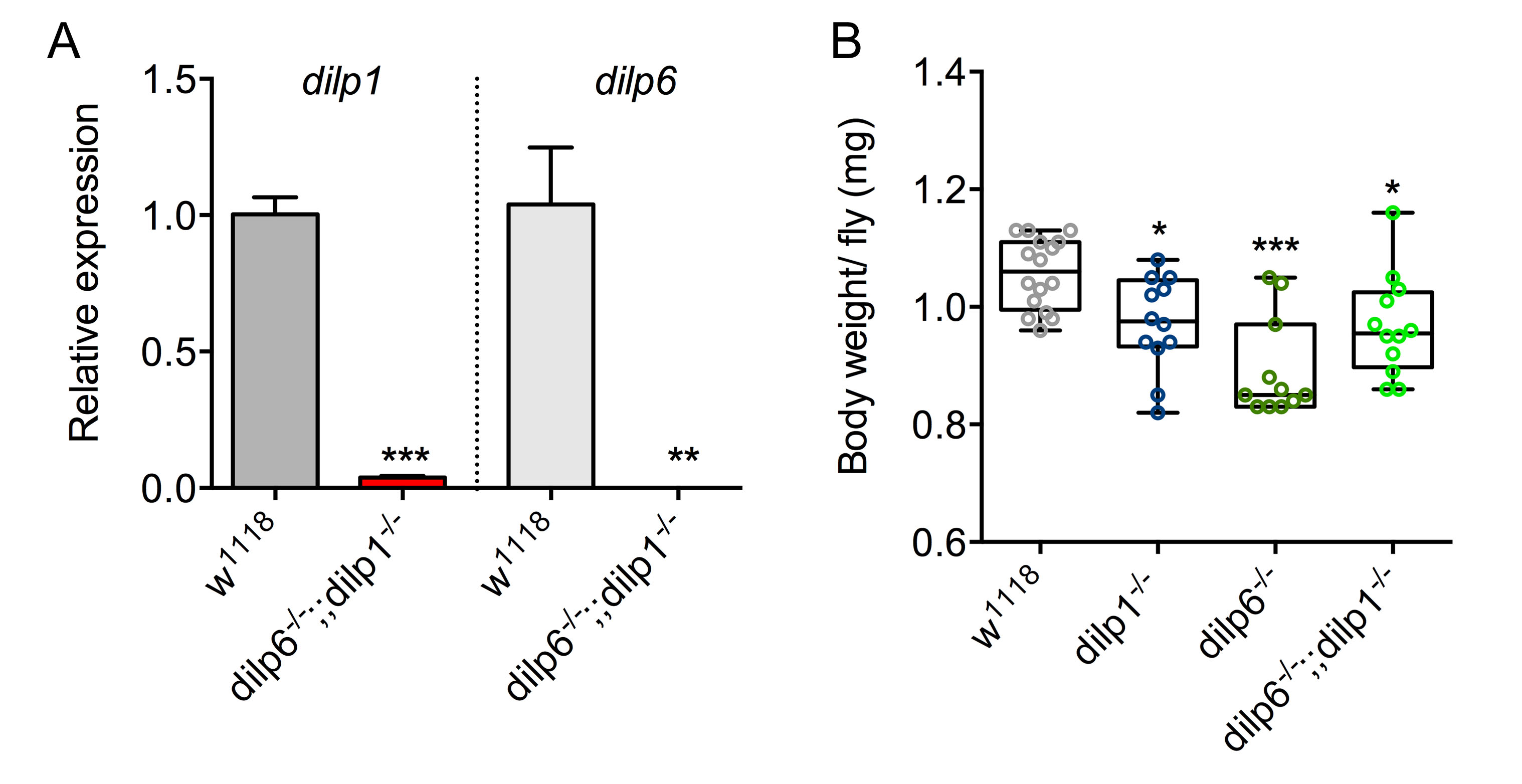

### S Fig. 3.jpg

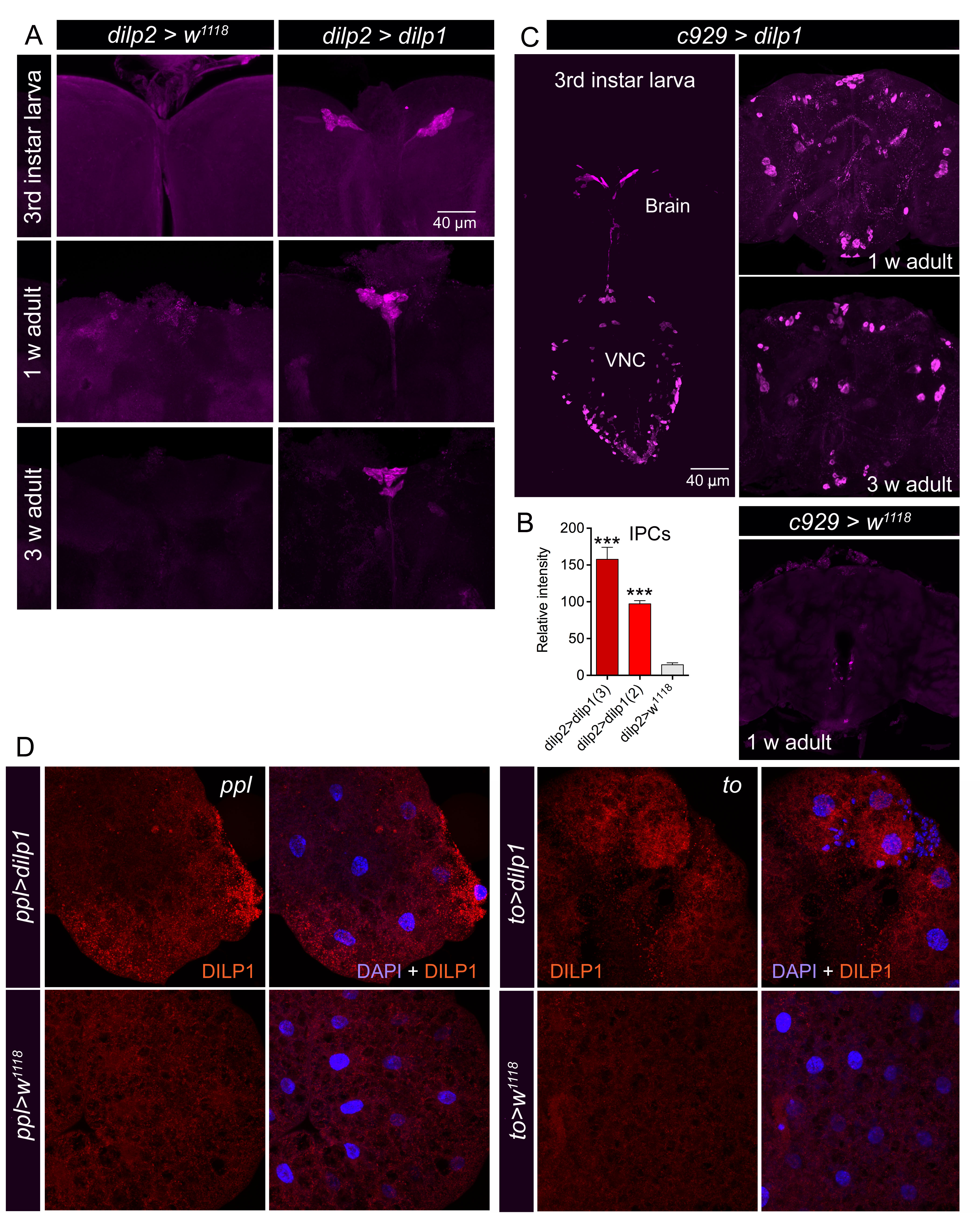

### S Fig. 4.jpg

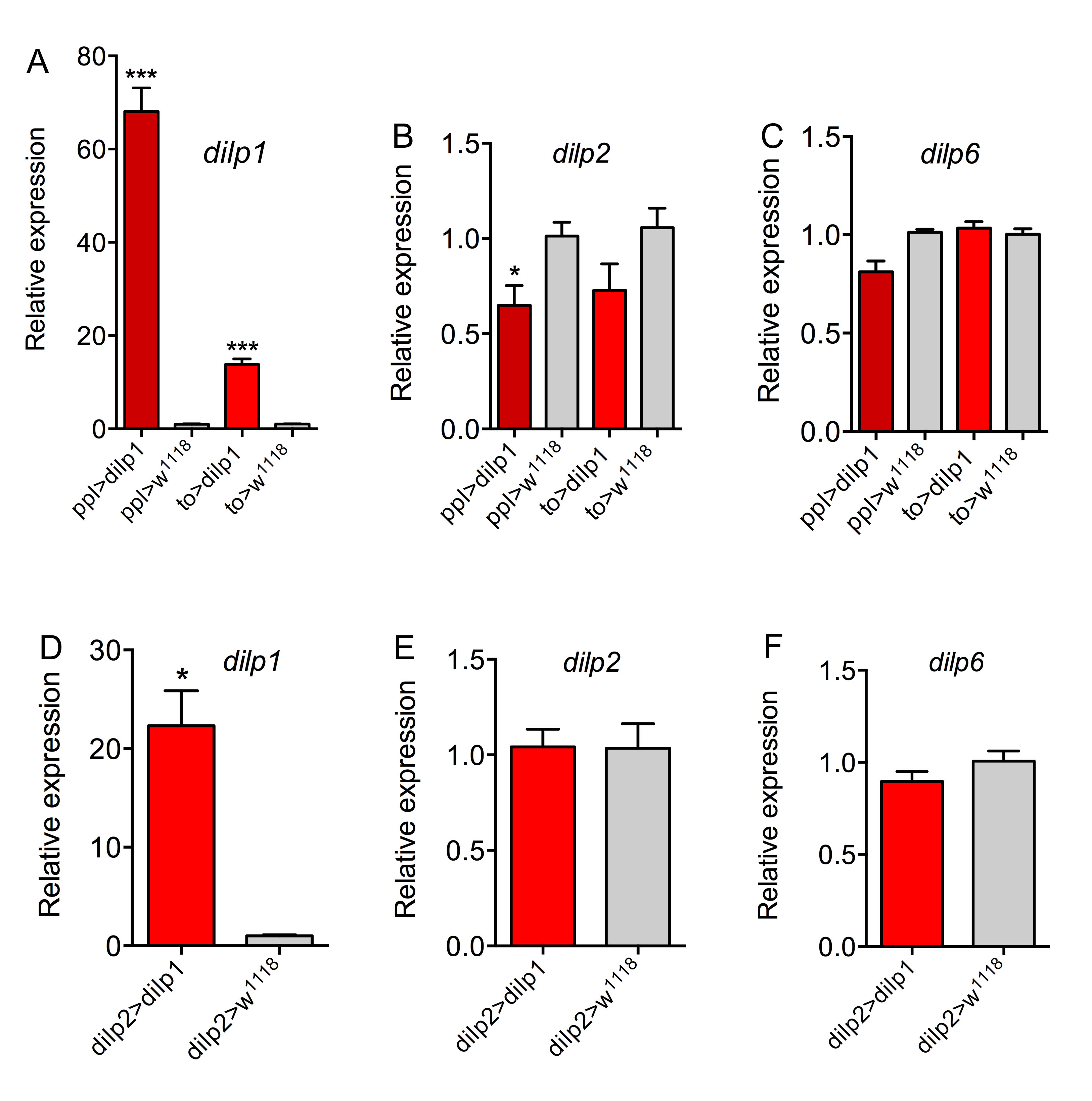

### S Fig. 5.jpg

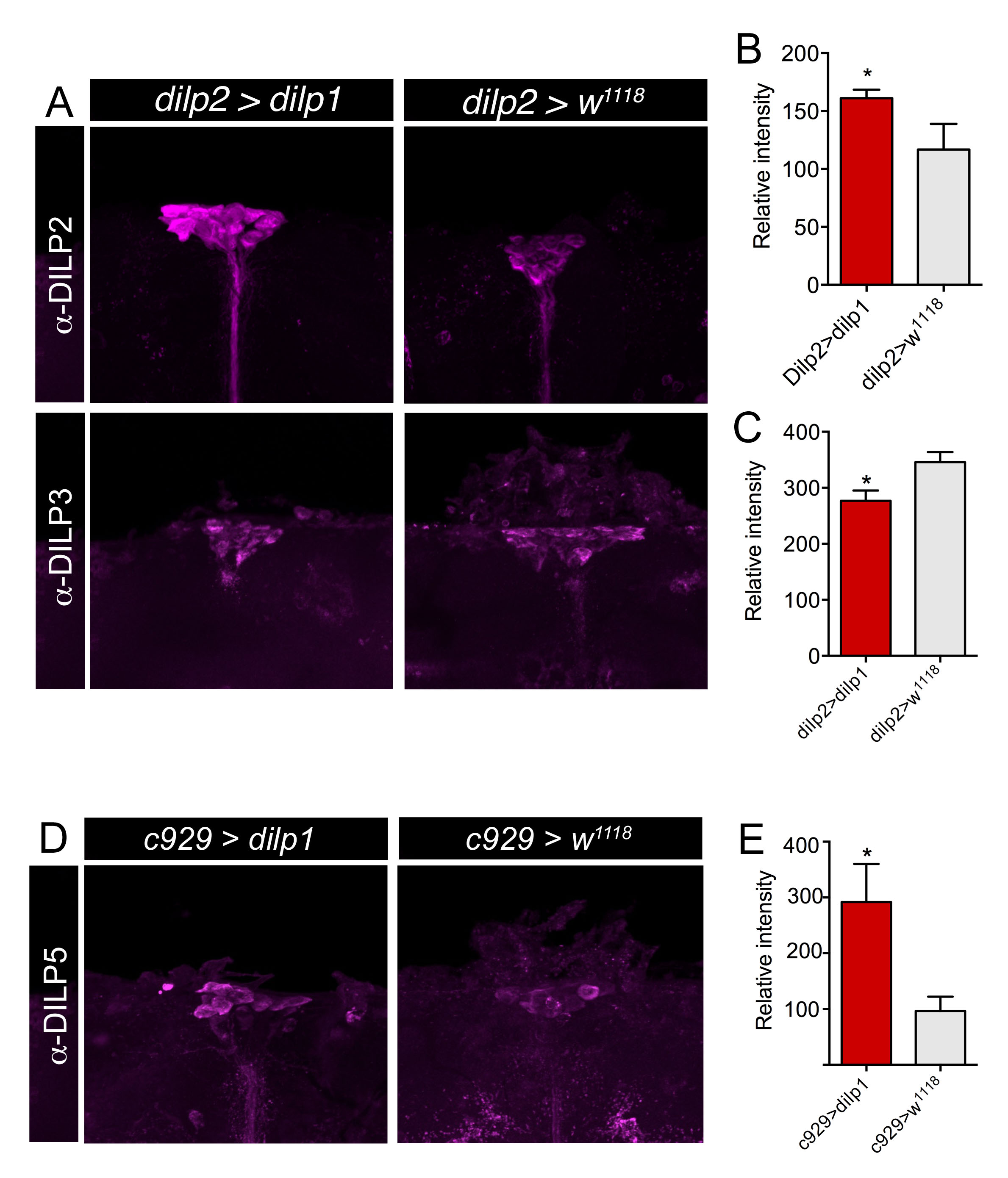

### S Fig. 6.jpg

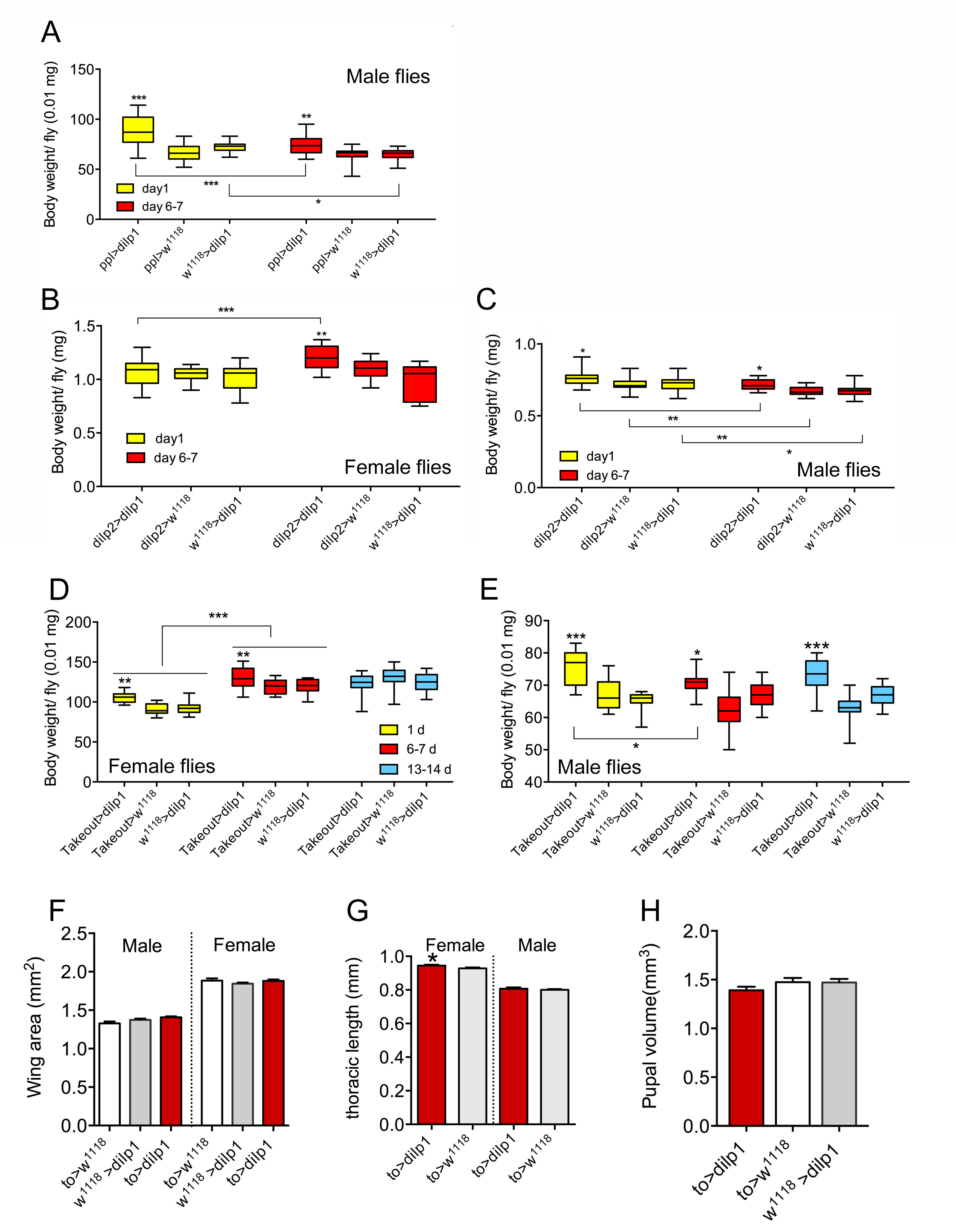

### S Fig. 7.jpg

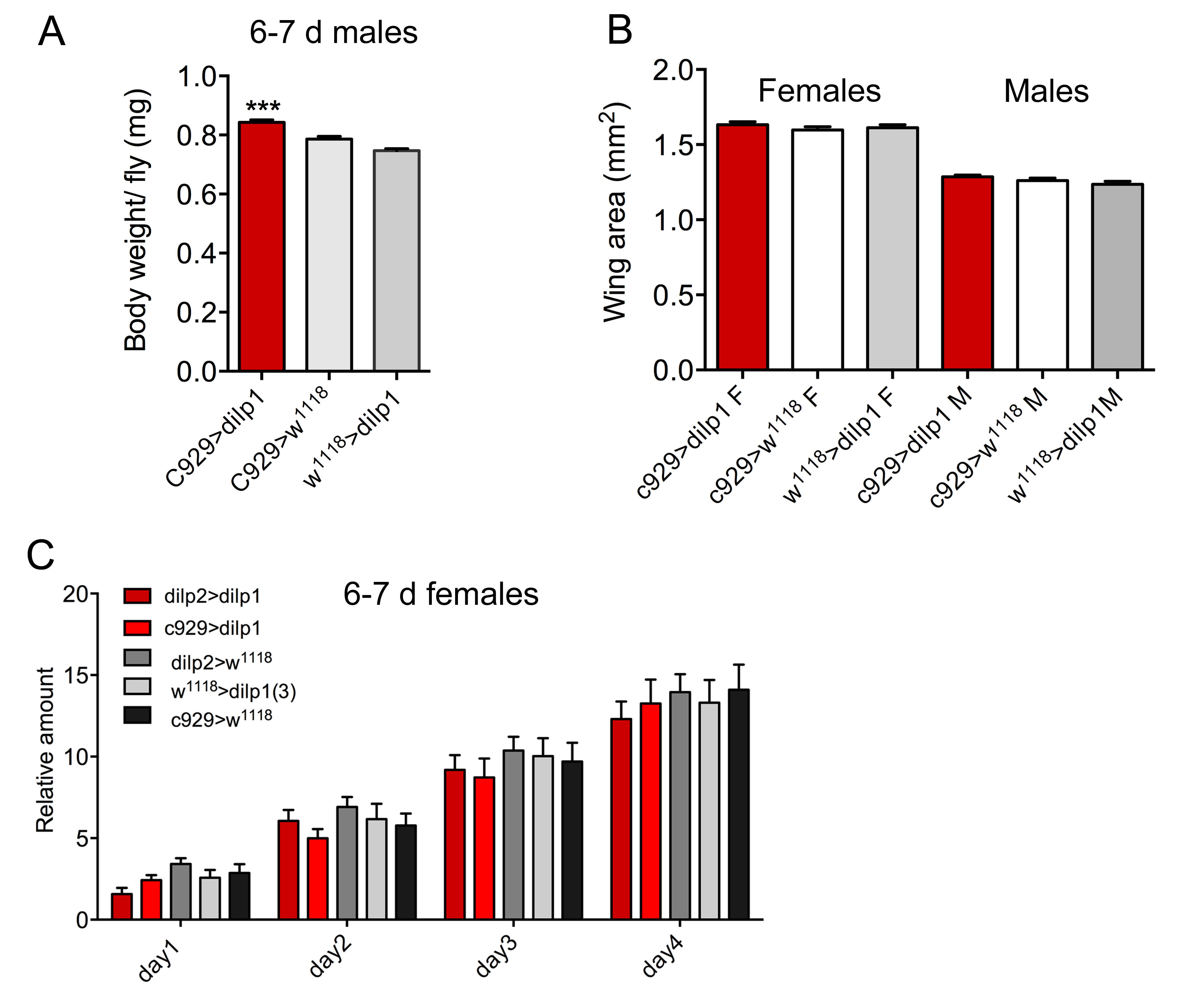

### S Fig. 8.jpg

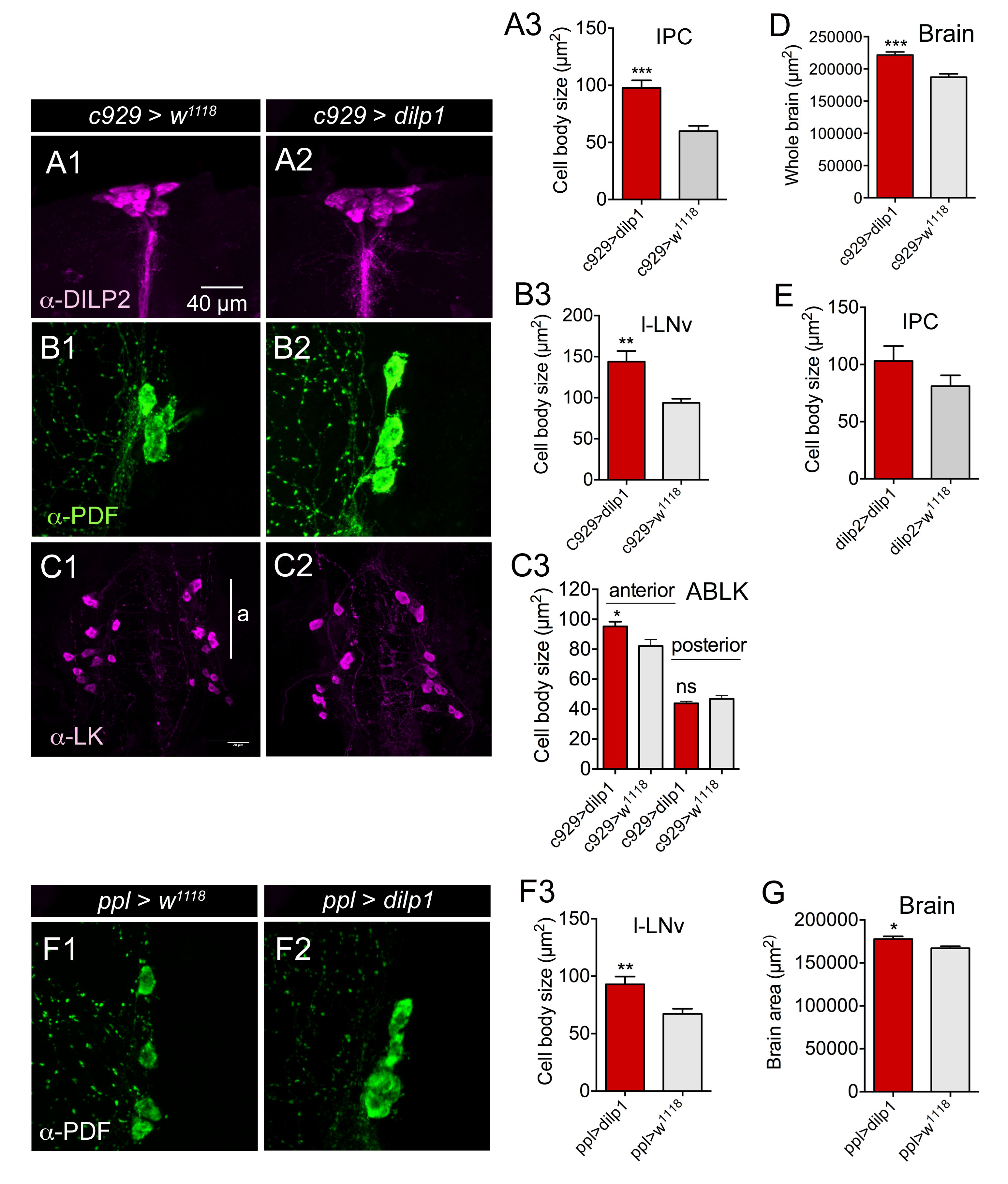

### S Fig. 9.jpg

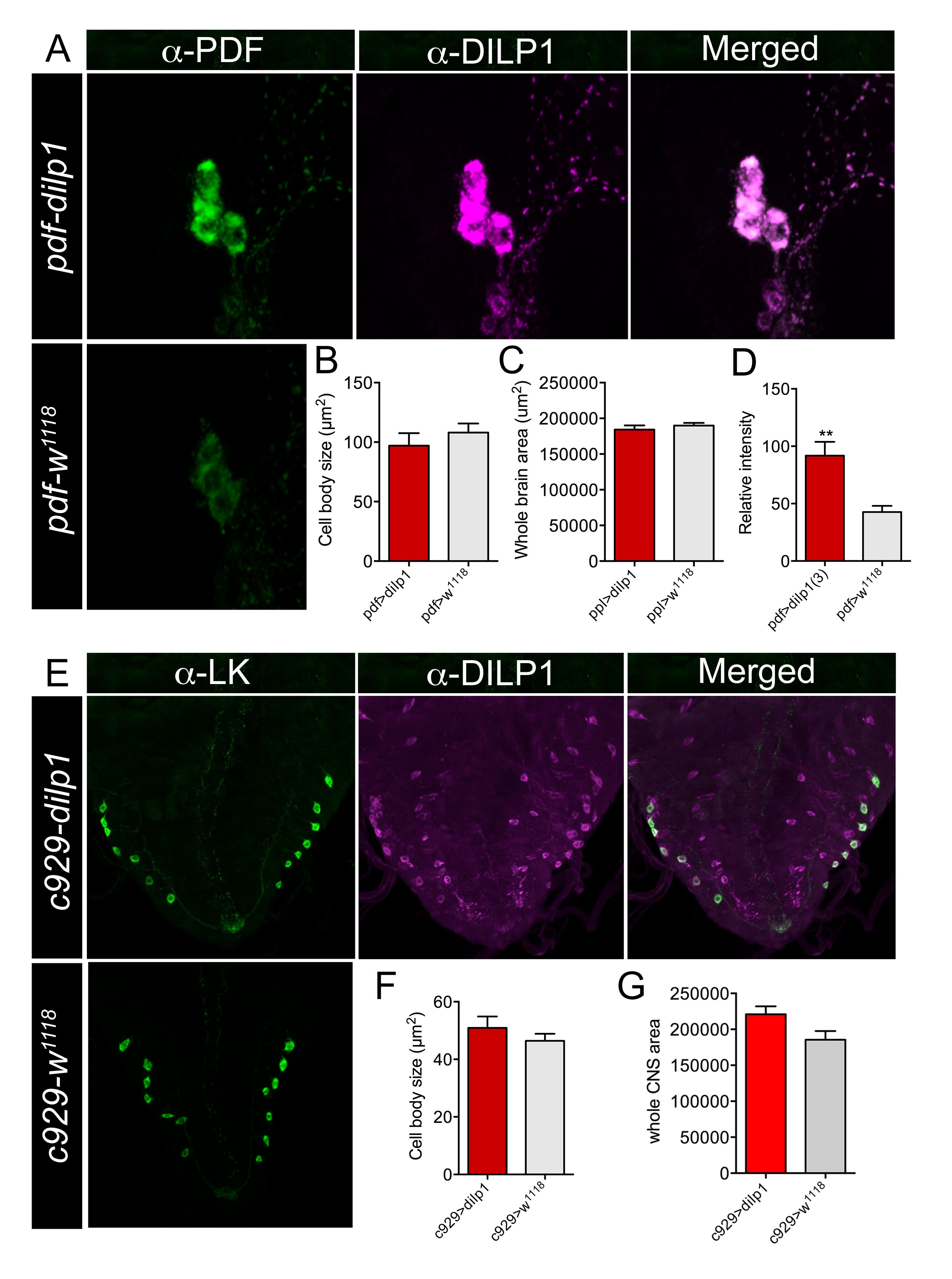

### S Fig. 10.jpg

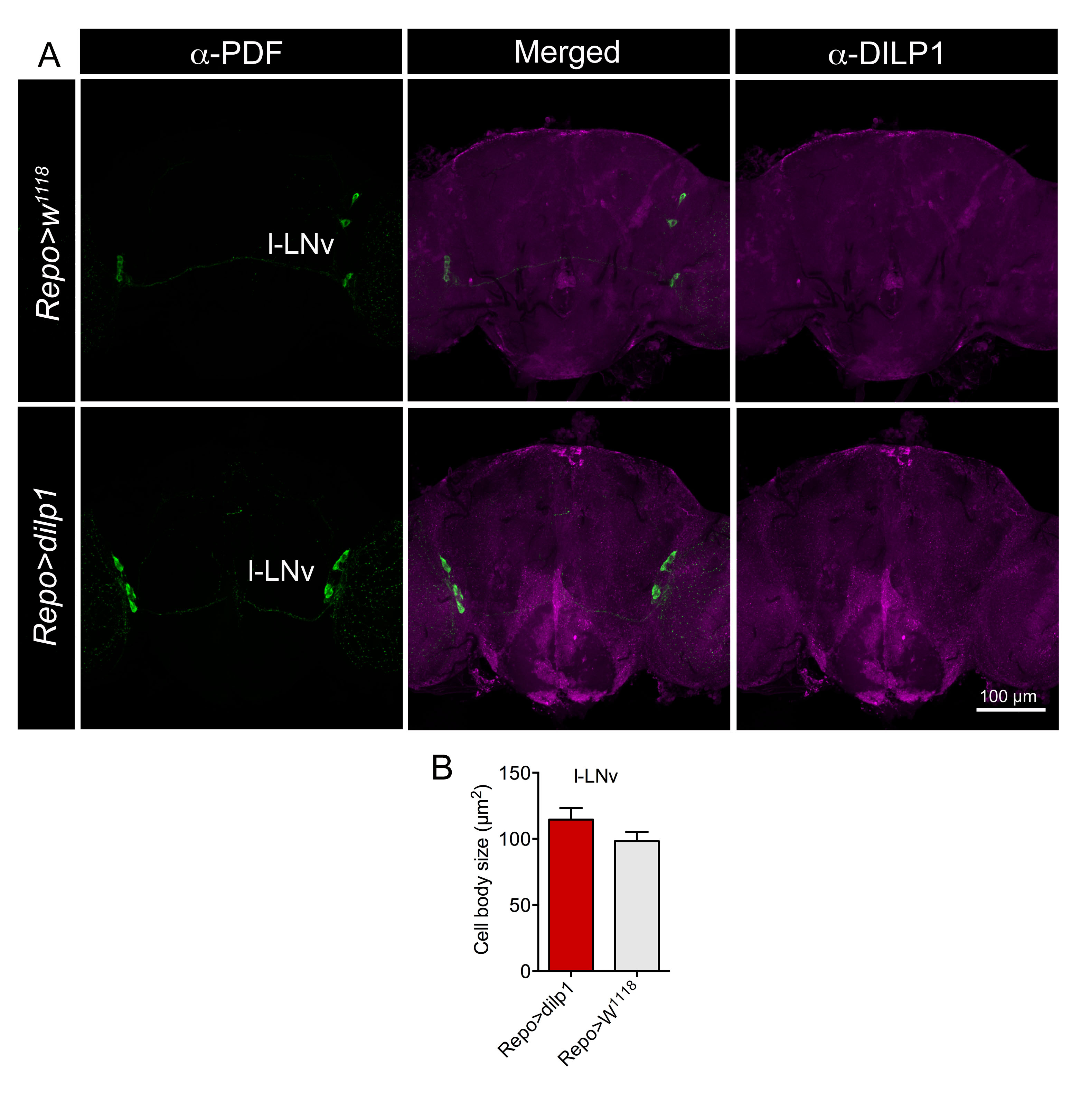

### S Fig. 11.jpg

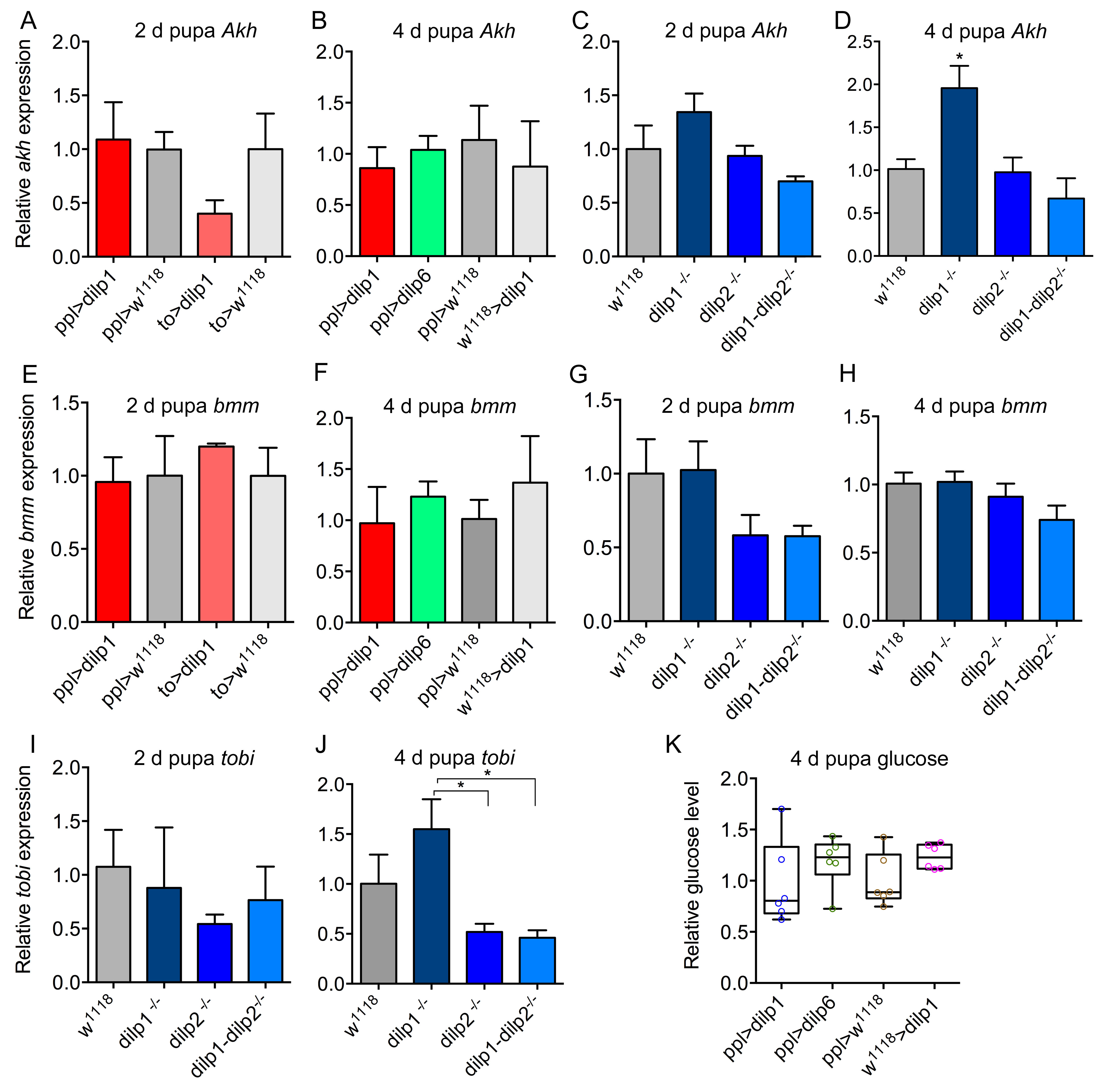

### S Fig. 12.jpg

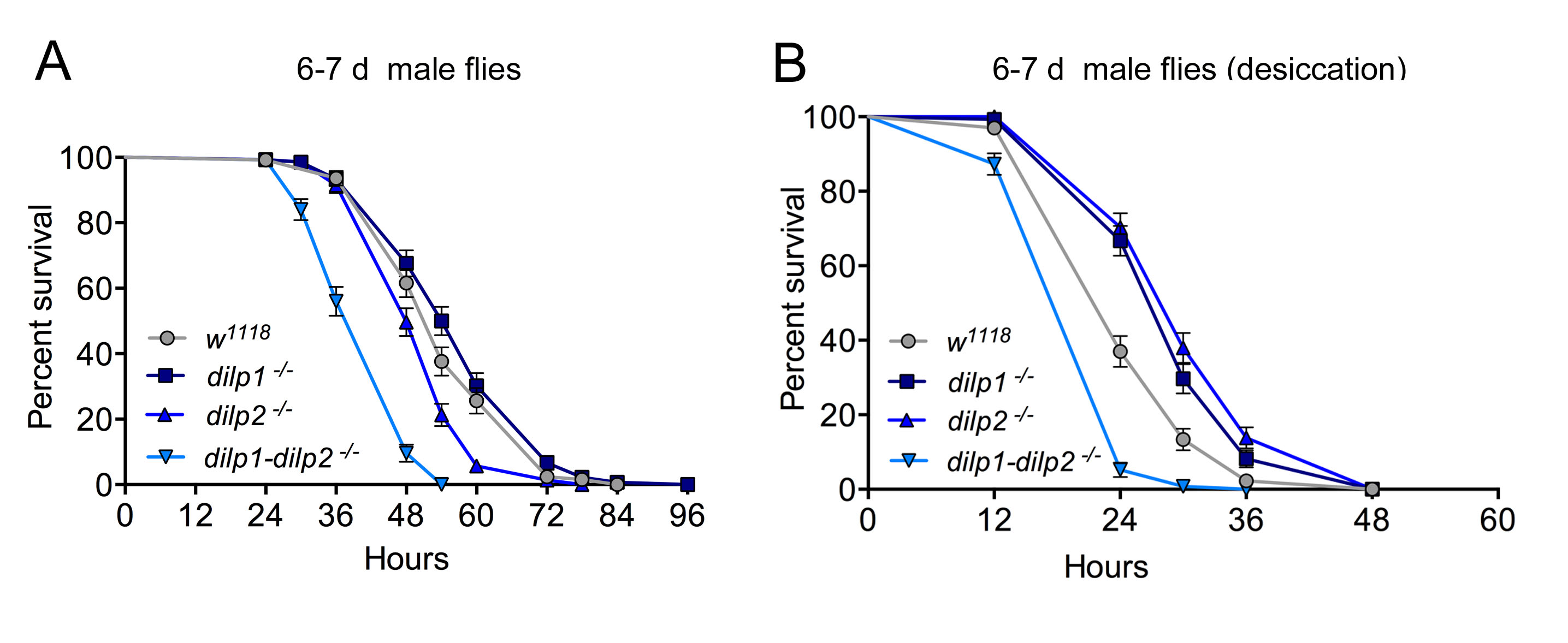

### S Fig. 13.jpg

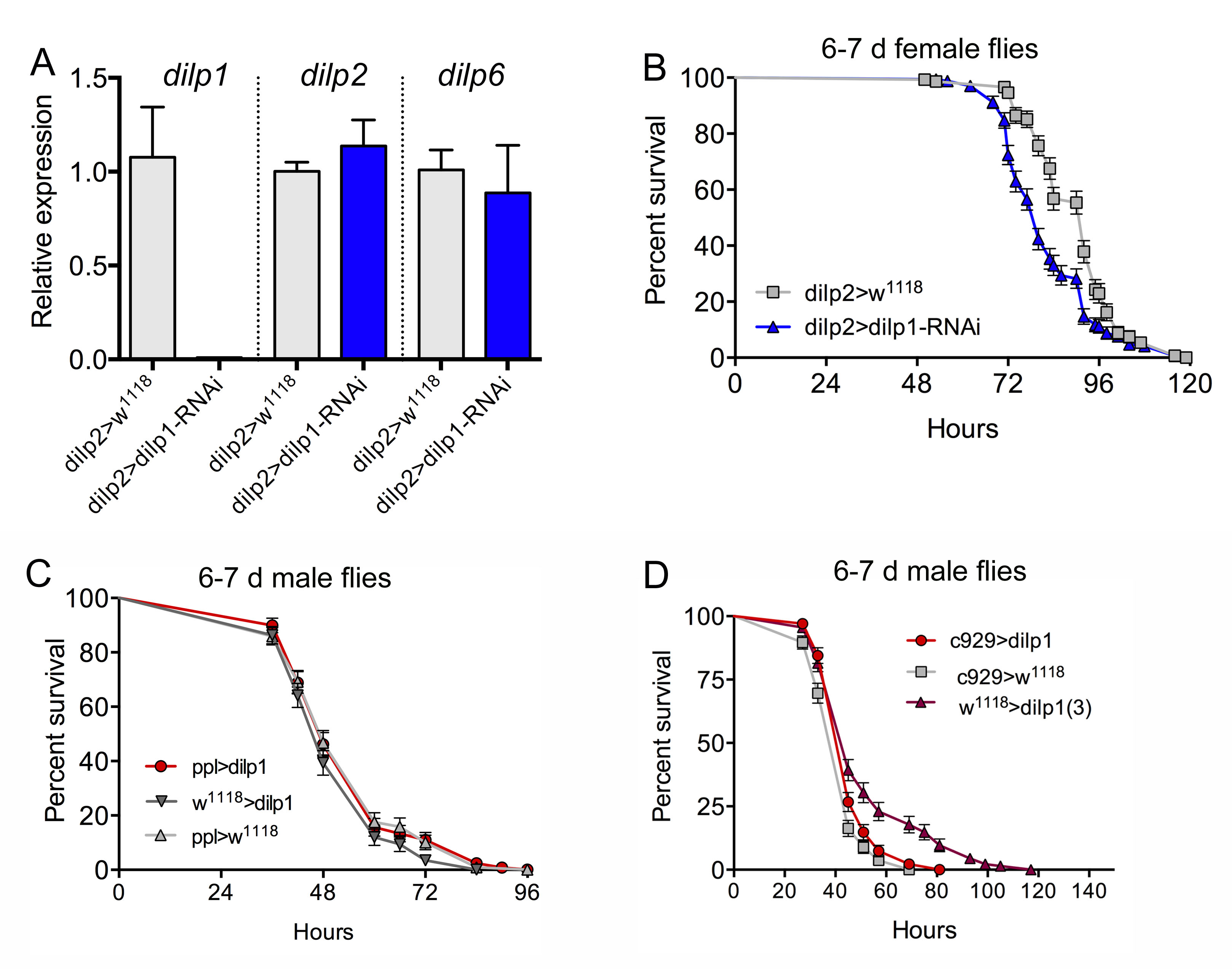
